# A cross-species study of sex chromosome dosage effects on mammalian brain anatomy

**DOI:** 10.1101/2022.08.30.505916

**Authors:** Elisa Guma, Antoine Beauchamp, Siyuan Liu, Elizabeth Levitis, Liv S. Clasen, Erin Torres, Jonathan Blumenthal, Francois Lalonde, Lily R. Qiu, Haley Hrncir, Allan MacKenzie-Graham, Xia Yang, Arthur P. Arnold, Jason P Lerch, Armin Raznahan

## Abstract

All eutherian mammals show chromosomal sex determination with contrasting sex chromosome dosages (SCDs) between males (XY) and females (XX). Studies in transgenic mice and humans with sex chromosome trisomy (SCT) have revealed direct SCD effects on regional mammalian brain anatomy, but we lack a formal test for cross-species conservation of these effects. Here, we develop a harmonized framework for comparative structural neuroimaging and apply this to systematically profile SCD effects on regional brain anatomy in both humans and mice by contrasting groups with SCT (XXY and XYY) vs. XY controls. We show that total brain size is substantially altered by SCT in humans (significantly decreased by XXY and increased by XYY), but not in mice. Controlling for global effects reveals robust and spatially convergent effects of XXY and XYY on regional brain volume in humans, but not mice. However, mice do show subtle effects of XXY and XYY on regional volume, although there is not a general spatial convergence in these effects within mice or between species. Notwithstanding this general lack of conservation in SCT effects, we detect several brain regions that show overlapping effects of XXY and XYY both within and between species (cerebellum, parietal, and orbitofrontal cortex) - thereby nominating high priority targets for future translational dissection of SCD effects on the mammalian brain. Our study introduces a generalizable framework for comparative neuroimaging in humans and mice and applies this to achieve a cross-species comparison of SCD effects on the mammalian brain through the lens of SCT.

**Highlights:**

- Parallel structural neuroimaging in humans and mice with sex chromosome trisomies
- Divergent X- and Y-chromosome effects on human brain size, but convergent effects on regional anatomy
- Muted impact of additional X or Y on mouse brain, but subtle regional effects evident
- Evidence for conserved cross-species impact of X and Y on fronto-parietal cortices and cerebellum

## 1. Introduction

Several independent lines of research provide evidence for sex chromosome dosage (SCD) effects on mammalian brain anatomy. In humans, sex chromosome aneuploidies (SCA) - a group of neurogenetic disorders characterized by carriage of an abnormal number of X- and/or Y-chromosomes - are associated with significant neuroanatomical alterations. More specifically, abnormal SCD influences both total volume and regional anatomical configuration of the human brain. Furthermore, neuroimaging studies of “four core genotype” mice - a model that allows for chromosome dosage and gonadal background to be uncoupled - have identified SCD effects wherein XX and XY groups differ in regional brain anatomy independent of gonadal context ^1,2^.

To date, we lack a formal comparison of SCD effects on the human and murine brain, which would be valuable for several reasons. First, it would provide a well-defined setting to develop new tools for the task of formally comparing neuroanatomical changes between species. To date, we lack analytic frameworks for asking if a variable of interests induces a spatially similar pattern of regional brain changes in humans and mice. Addressing this gap could accelerate comparative neuroscience more generally. Second, formally assessing homology of SCD effects between humans and mice is crucial for specifying the ways in which mouse models of SCD could serve to model mechanistic aspects of SCA in humans. Third, the unusual synteny of X-chromosome between humans and mice ^3^ increases the translational potential for exploring cross-species homology in gene dosage alterations. Indeed, there are several obstacles to mechanistic dissection of these effects in humans, including, but not limited to, the inaccessibility of the human brain, the difficulty in uncoupling SCD from gonadal effects, and environmental variation. The Sex Chromosome Trisomy mouse model produces XXY, XYY, XY, and XX mice with either a gonadal male (testes) or female (ovaries) background ^4^. Importantly, the model has been shown to recapitulate metabolic and motor features found in humans with sex chromosome trisomies (SCTs) ^4^, however, the neuroanatomical effects have neither been characterized in this model nor compared to human SCT effects. Fourth, although SCTs are defined by an abnormal number of sex chromosomes, they provide a rare means of asking if foundational sex-biased biological factors - namely X- and Y-chromosome count - are associated with similar brain features in two different eutherian mammals.

Here, we use comparative neuroimaging in humans and murine cohorts with the same SCT variations, namely XXY and XYY, relative to XY controls, to assess the effect of an added X- or Y-chromosome on structural neuroimaging-derived global and regional brain volume. We evaluate the degree of spatial convergence between the effect of an added X- or Y-chromosome in each species. Finally, for a set of homologous human-mouse brain regions we assess the similarity of effects for an added X- or Y-chromosome across species. Our approach harnesses recent advances in neuroimaging methods leveraging high-field magnets that have allowed researchers to study the brain of rodents in a noninvasive way, acquiring comparable signal to that acquired in human neuroimaging studies ^5–7^. Our focus on structural MRI derived phenotypes is motivated by (1) the known effects of sex and SCD on brain volume from both human and mouse neuroimaging studies ^1,2,8,9^ and (2) the advantages of using brain volume over behavioural measures for cross-species comparisons given the complexities of mapping human behaviour in nonhuman animals ^10^. Of note, although this current study applies a workflow for comparative neuroimaging to probe SCD effects, the methodological approaches we introduce can be generalized to aid comparative analysis of many other influences on brain anatomy.

## 2. Results

### 2.1 Comparing SCT effects on global brain volume in humans and mice

We first examined the effects of added X- or Y-chromosomes on total tissue volume (TTV) in humans (_H) and mice (_M). Each human SCT group (XXY_H, and XYY_H) had an independent control group (XY_H) (due to a scanner software upgrade between collection of XXY and XYY data), while the two murine aneuploidy groups (XXY_M, XYY_M) shared a single control group (XY_M). In humans volume was significantly decreased in XXY_H (−4.3%, 95% confidence interval (CI) [−5.6%,−3.0%]; t=−6.429, β=−0.944, p=1.14e^−09^), and non-significantly increased in XYY_H (+1.5%, 95% CI [-0.5%,+3.4%]; t=1.561, β=0.423, p=0.123) (**Fig 1AC**). In contrast, SCT had a negligible effect on TTV in mice, with slightly smaller volumes in both aneuploidy groups as compared to controls (XXY_M vs XY_M: −0.6%, 95% CI [−2.1%,+0.9%]; t= −0.789, β= −0.285, p= 0.436; XYY_M vs XY_M: −0.7%, 95% CI [−2.3%,+0.9%]; t=−0.860, β=−0.319, p=0.395) (**Fig 1BD**).

**Figure 1.**
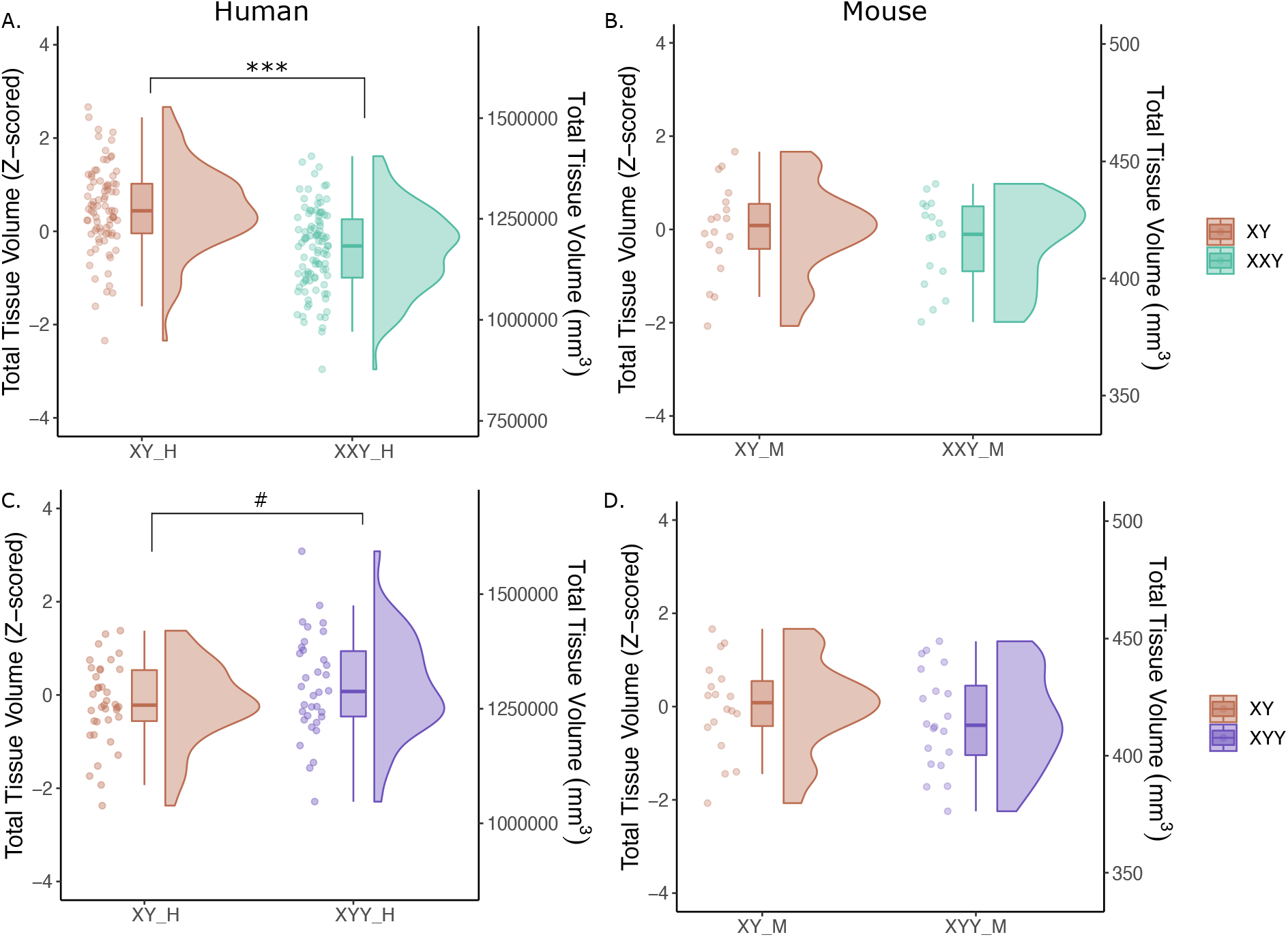
Effects of added X- or Y-chromosome on total tissue volume (TTV) in humans and mice. Distributions of TTV are shown for the effects of a supernumerary X- and Y-chromosome (**A,B** and **C,D** respectively) in humans (**A,C**) and mice (**B,D**). Data are represented using individual points, boxplot, and half-violin plot (raincloud plot). ***p<0.0001. # p=0.123; range of standardized effect sizes are as follows: XXY_H group β=−1.10 to +0.91; XYY_H β=−1.40 to +1.31; XXY_M group β=−0.78 to 0.61; XYY_M group β=−0.59 to 0.47.

### 2.2 Comparing SCT effects on regional brain volume in humans and mice

Next, we examined the effects of added X- or Y-chromosomes on regional gray matter volumes in each species. Human cortical volumes were derived from the Glasser-HCP parcellation (n=360) ^11^, a highly detailed, multi-modally informed cortical atlas, while subcortical human volumes were derived from the FreeSurfer parcellation (n=40) ^12^. Murine regional brain volumes were derived from segmentation using a previously published, highly detailed MRI-atlas of the adult mouse brain (n=355 regions) ^13–16^. In each species, regional volumes for each individual were scaled relative to the mean and standard deviation of their respective XY controls. Given the prominent effect of SCT on TTV in humans, but not mice, we controlled for TTV when estimating effects of SCT on regional brain volume in each species to allow for a more focused test of cross-species similarity of SCT effects on local rather than global anatomy. We also included age as a covariate when modeling SCT effects on regional brain volume in humans (but not mice as ages were the same for all).

In humans, addition of a Y-chromosome induced a wider effect size range of regional brain volume changes than addition of an X-chromosome (**Figure 2A**, Levene’s test for variance difference in effect sizes: F(1,754)=34.214, p=7.349e^−09^), whereas the distribution of effect sizes for mice with added X- or Y-chromosomes was not different (**Figure 2B**, Levene’s test, F(1,906)=0.224, p=0.636). For both XXY and XYY contrasts, the distribution of SCT effect sizes on regional brain volume was more variable in humans than mice (XXY: F(1,831)=14.091, p=0.0002; XYY: F(1,831)=109.83, p=2.263-16).

**Figure 2.**
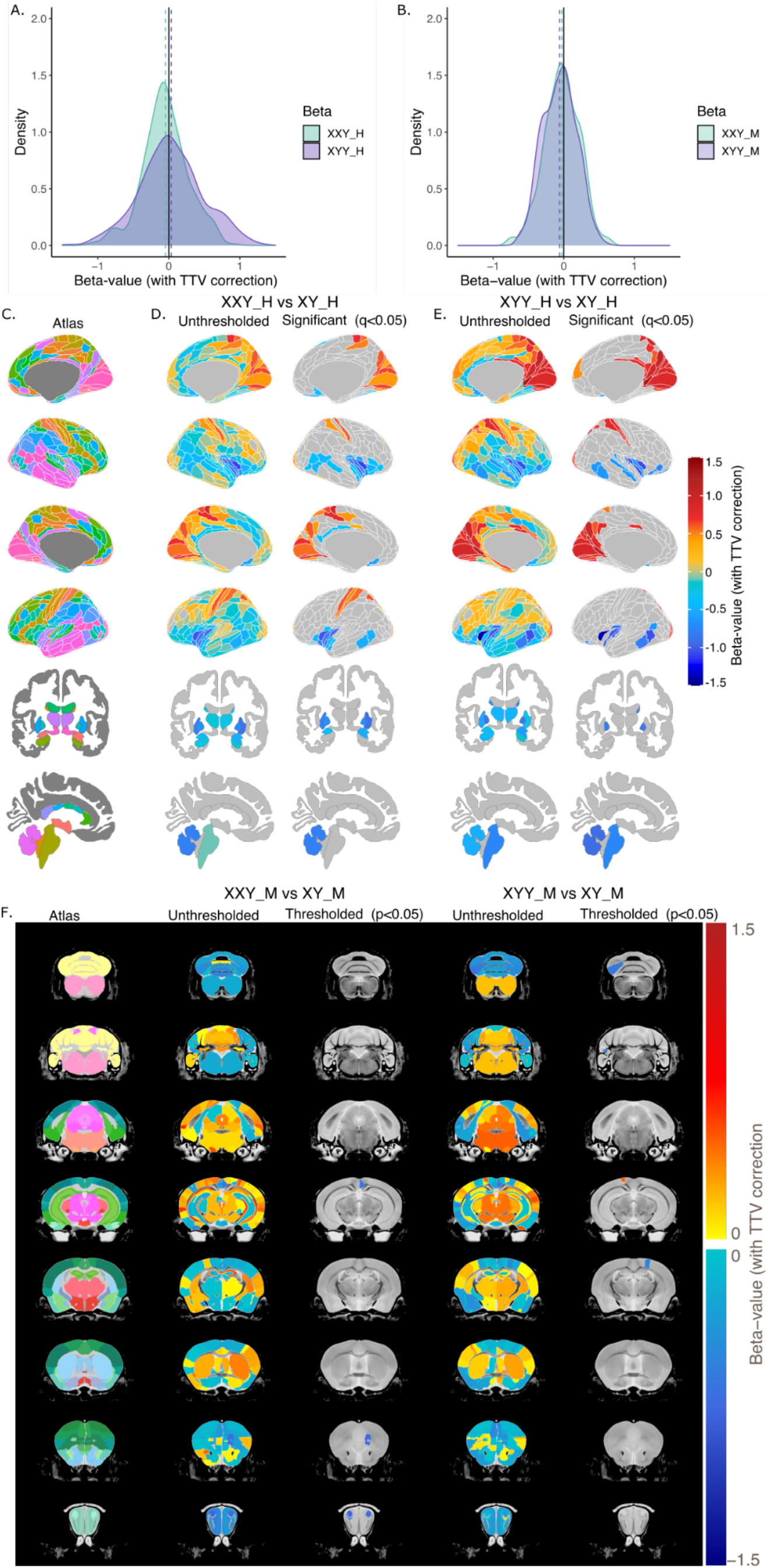
Effect of added X- or Y- chromosomes on regional brain volume in humans and mice, with total tissue volume (TTV) correction. Distribution of beta values for the effect of sex chromosome trisomy are displayed for humans (**A**) and for mice (**B**). **C.** Representative views of the cortical Glasser atlas for lateral and medial slices of the left and right hemispheres, and subcortical FreeSurfer atlas with both axial and sagittal views. Unthresholded (left) and significant (q<0.05; right) beta values for the effect of added X (**D**) and added Y (**E**) in the human brain. **F.** Mouse brain coronal slices with the representative atlas on the left, followed by unthresholded, then thresholded (p<0.05) beta values for the effect of added X, followed by unthresholded, then thresholded (p<0.05) effects of added Y. Human = _Y; mouse = _M.

Humans showed several statistically-significant effects of SCT on regional cortical and subcortical volume after correction for multiple comparisons (using the False Discovery Rate [FDR] correction ^17,18^ with q=0.05 and corresponding t-value thresholds of XXY_H: t=2.678, XYY_H: t=−2.528). We observed highly similar effects of XXY_H and XYY_H on regional brain anatomy. In humans, both SCTs induced statistically significant volume increases in the bilateral posterior parietal cortex (XXY_H: 5m, 5mv, 5L, 7AL, and left 7PL, 7m, 7Am; XYY_H: 5L, R5m, 7AL, R7Am, 7m), visual cortex (XXY_H: V1, V2, and left V3, V7; XYY_H: V1, V2, V3, L V3A, V4, V6, L V6A, V7, and prostriate cortex), dorsal visual transition area, parieto-occipital sulcus, intraparietal area, primary somatosensory cortex (XXY_H: area 1, 2, 3b; XYY_H: area 1, 2), and cingulate cortex (XXY_H: 24dd; XYY_H: v23ab, p24pr, 31pd). In both XXY_H and XYY_H, volume decreases were seen in the insular cortex (XXY_H: MI, Ig, Pol1 & 2, AI, AAIC, R 52; XYY_H: Pol1, MI, 52, R Pol2, L Ig, L AVI), temporal cortex (XXY_H: TPOJ1 & 2, TA2, STGa, STSdp, and bilateral PHT; XYY_H: TE1p, PHT, R STSda, L TPOJ2), orbital part of inferior frontal gyrus (area 47), orbitofrontal cortex (OFC), fusiform face complex (bilateral in XXY_H and only right hemisphere in XYY_H), piriform cortex (bilateral in XXY_H, and right hemisphere in XYY_H), bilateral pallidum, and cerebellar cortex were observed in both human SCT groups. Despite this general picture of spatial convergence between significant XXY_H and XYY_H effects on regional brain volume, a few dissociations were seen. For example, XXY_H individuals displayed greater increases in the posterior parietal cortex, particularly in the left hemisphere, while XYY_H increases in the visual cortex were more prominent. Additional statistically significant volume decreases were observed in the bilateral putamen, pallidum, and amygdala for the XXY_H group but not the XYY_H group (**Figure 2DE**).

Mice lacked statistically significant effects of SCT on regional brain volumes after correction for multiple comparisons (using the False Discovery Rate [FDR] correction with q=0.05). At a relaxed threshold (uncorrected p<0.05) we observed discordant effects due to XXY and XYY, wherein XXY_M had volume decreases in the accessory olfactory bulb and ventral retrosplenial area, specifically, the right cingulate cortex (area 29C), the right orbital area of XXY_M, and in the right primary somatosensory area and the left paramedian lobule of the cerebellum, and volume increases in the perirhinal area, while XYY_M had volume decreases in the primary somatosensory cortex and paramedian lobule of the cerebellum (**Figure 2F**).

The SCT mouse model produces sex chromosome variations (e.g., XY, XXY, XYY) with either testes or ovaries (because of the presence or absence of the *Sry* transgene) ^4^. Therefore, in mice only, we were able to further test the stability of SCT effects on regional brain volume on an ovarian background (in addition to the testicular background necessarily used for our primary comparison of SCT effects in humans and mice). This supplementary test in mice revealed a moderately strong cross-regional correlation in the volumetric effects of XYY between ovarian and testicular groups (r=0.557) and a weaker cross-regional correlation in XXY effects (r=0.216) (**Supplementary Figure 2**). Combining testicular and ovarian groups to model the main effects of XXY and XYY with increased power in the largest murine sample available (controlling for gonadal type and TTV) revealed statistically significant decreases at 10% FDR in the olfactory bulbs of both XXY and XYY mice relative to XY (glomerular layer of the accessory olfactory bulb for XXY and right plexiform layer of the main olfactory bulb for XYY).

Repeating the above analyses without TTV correction (details in **Supplement 1.1 & Supplementary Figure 1**) did not reveal any statistically significant effects of SCT on regional brain volume in mice, and modified findings in humans such that significant brain-wide volume differences were observed in the XXY_H group (driven by smaller TTV), while effects in XYY_H were similar to those described above (details in **Supplement 1.1 & Supplementary Figure 1**). See **Supplementary Tables 2-3** for human model outputs and **Supplementary Tables 4-5** for mouse model outputs (with and without TTV correction).

### 2.3 Testing for convergent SCT effects in each species

The above results suggested that after correction for TTV effects, humans show statistically significant and largely convergent effects of XXY and XYY on regional brain volume, whereas mice do not show statistically significant effects of SCT on regional brain volume. We therefore developed a resampling-based approach to formally test for spatial convergence of more subtle SCT effects in mice while ensuring independence of XY controls between murine SCT groups (so as to mirror the independence of XY controls for SCT groups in humans). Specifically, for each round of this analysis, we generated two independent XY control groups by repeatedly splitting the XY_M control group (n=18) in half, sample A (n=9) and sample B (n=9), sampling without replacement. Regional effect sizes were estimated for XXY and XYY as compared to their respective XY control groups (sample A for XXY and sample B for XYY), and the cross-ROI correlation in these effect sizes was computed. This procedure was repeated 1000 times to (i) yield a distribution of estimates for the cross-ROI similarity in effect sizes of XXY and XYY on murine brain volume, and (ii) identify any ROIs that showed a congruent direction of volume change in XXY and XYY across at least 95% of these iterations. We took any such regions to be instances of sub-threshold convergence in the effects of XXY and XYY on regional murine brain volume.

The correlation of standardized effect sizes for each of the bootstrap resampled splits was low (cross-ROI mean r=−0.053), with the full distribution of these correlations across all 1000 splits displayed in **Supplementary Figure 4A**. The cross-ROI correlation in effect sizes for XXY and XYY in humans was high (**Fig 3A**, r=0.734), while in mice there did not appear to be a global alignment between the spatial pattern of XXY and XYY effects on regional brain volume (cross-ROI effect size correlation for a representative murine split, **Fig 3B**, r=0.073). However, despite this lack of globally convergent XXY and XYY effects in mice, we identified several regions which showed convergent volume changes between XXY_M and XYY_M groups above chance levels (p<0.05). These regions allow a comparison with the regions of convergent statistically significant effects of XXY and XYY on regional volume in humans, which include increased volume in the posterior occipito-parietal regions and ventral anterior cingulate cortex, along with convergent volume decreases in insular, lateral temporal, orbitofrontal, piriform, fusiform face, pallidal, and cerebellar regions (**Fig 3C**). In mice (**Fig 3D**), we observed consistently overlapping volume increases in XXY_M and XYY_M vs. XY_M controls in the medial parietal association cortex (also increased in human SCT **Fig 3E,I**), right dorsal pallidum (decreased in human SCT **Fig 3F,J**), right CA1 and subiculum, right perirhinal area, and the periaqueductal gray. Conversely, consistent volume decreases with SCT in mice were seen in the olfactory bulbs (glomerular, mitral, and plexiform layers) as well as in the right cingulate cortex (29c) (also decreased in human SCT **Fig 3G,K**), right primary somatosensory cortex (increased in human SCT), right orbital area (also decreased in human SCT), and the left paramedian lobule of the cerebellum (also decreased in human SCT **Fig 3H,L**). We provide selected regional volume plots for SCT and XY groups from both humans (**Fig 3E-H**) and mice (**Fig 3I-J**) to illustrate the convergent and divergent neuroanatomical effects of SCT across species and highlight cross-species similarities and dissimilarities (**Fig 4**).

**Figure 3.**
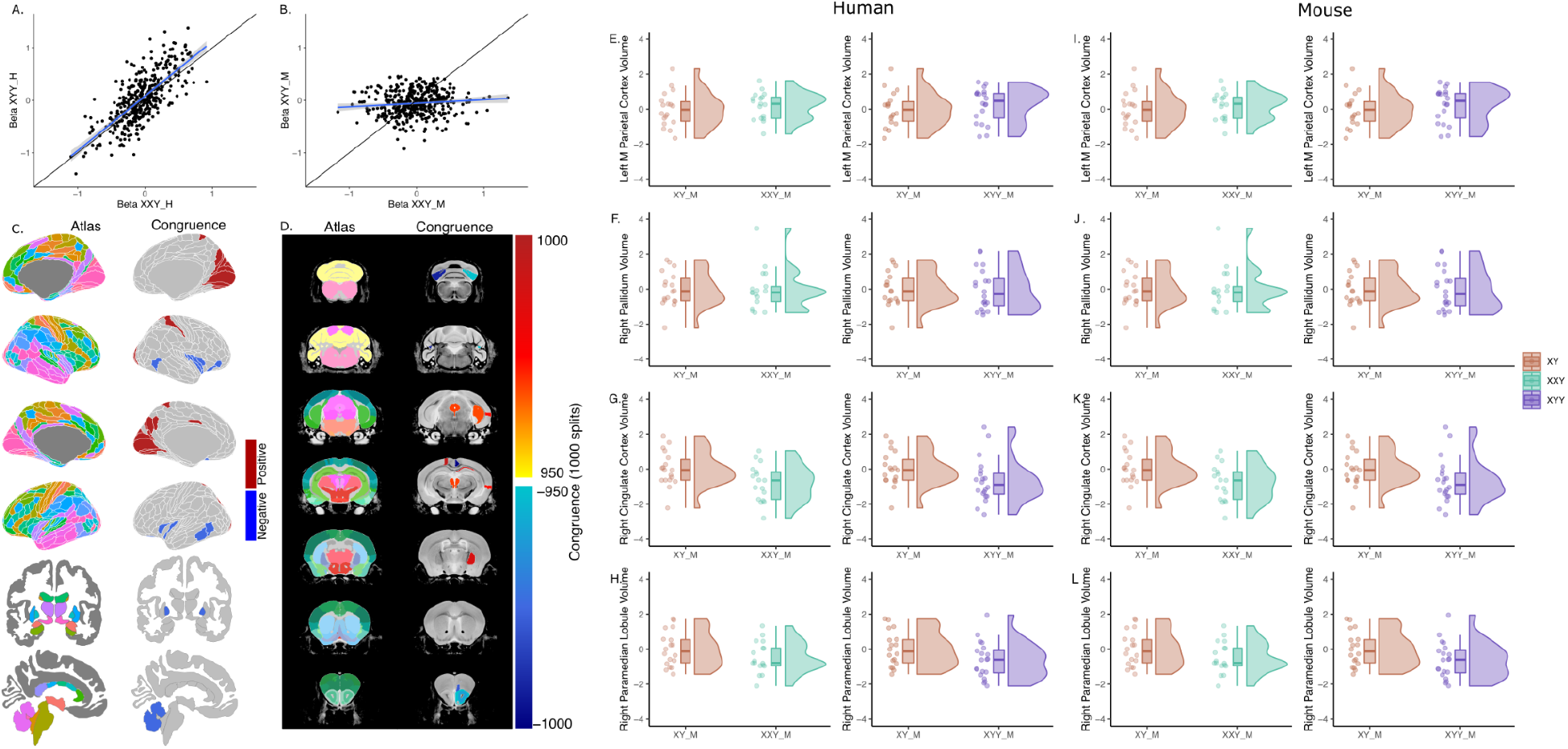
Spatial convergence between added X- or Y- chromosomes on the human and mouse brain. Standardized effect sizes for the effect of an added X- or added Y-chromosome are highly correlated across brain regions in humans (**A**), but not in mice [as illustrated by a representative split from the 1000 bootstraps (split 34; **B**)]. **C.** Human brain regions whose volume were either convergently increased (red; n=22) or consistently decreased (blue; n=18) in both aneuploidy groups. **D**. Mouse brain regions whose volume was consistently increased (red colours) or decreased (blue colours) in 95% (950/1000 splits) of the XY_M control splits in both aneuploidy groups based on bootstrap analysis. Human anterior cingulate cortex (**E**) and parietal cortex (**F**) volume were convergently increased in both XXY_H and XYY_H, and cerebellum (**G**) and pallidum (**H**) volume were convergently decreased in both XXY_H and XYY_H. Mouse cingulate cortex (**I**), parietal cortex (**J**), cerebellum (**K)** volume was consistently and convergently decreased in both XXY_M and XYY_M, and pallidum (**L**) volume was consistently and convergently increased in both XXY_M and XYY_M. All plots use the beta values for the effect of aneuploidy with total brain volume correction. For all boxplots, volumes are standardized (z-scored relative to XY controls). Human = _Y; mouse = _M.

**Figure 4.**
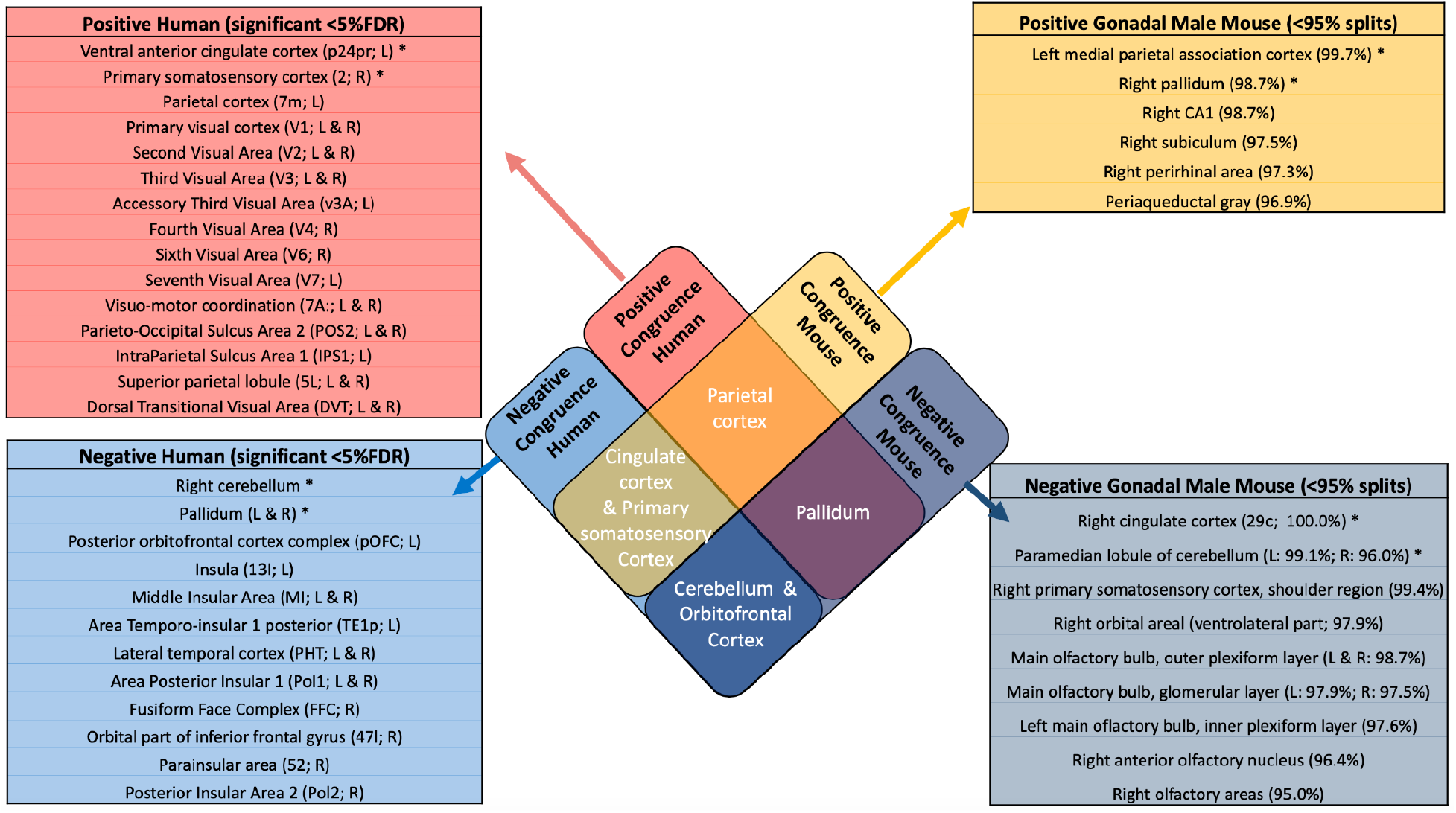
Convergently increased or decreased regions in humans and mice. Venn diagram of overlapping ROIs highlighting the regions congruently (for both XXY and XYY) increased or decreased in humans and mice. Brain regions that were statistically significantly impacted by both XXY and XYY in humans alone (<5%FDR) are listed in the red (increased volume) and light blue (decreased volume) boxes on the left. Brain regions that showed significantly convergent impacts of XXY and XYY (based on bootstrap resampling) in mice alone are listed in the yellow (increased volume) and dark blue (decreased volume) boxes. Intersection cells list regions showing convergent XXY and XYY effects in both species: orange - regions that are increased by XXY and XYY in both species; navy blue - regions that are decreased by XXY and XYY in both species; green - regions that are increased by XXY and XYY in mice, but decreased in humans; and, purple - regions that are increases by XXY and XYY in humans but decreased in mice. Human = _Y; mouse = _M.

Repeating the above tests for spatial convergence without covarying for TTV retained the strong spatial alignment between regional XXY and XYY effects in humans (cross-ROI effect size correlation r=0.731) and the lack of such an alignment in mice (cross-ROI correlation r= 0.024). Regions showing a sub-threshold overlap in XXY and XYY-induced volume decreases in mice when not covarying for TTV included the cerebellum (paramedian lobule and crus 2), the cortex (cingulate cortex area 29c, somatosensory cortex, orbital cortex, and prelimbic cortex), the subcortex (medial amygdalar nucleus), the brainstem (right pontine reticular nucleus), and the olfactory bulbs. A number of these regions align with those observed in the TTV corrected analyses (cerebellum, somatosensory, cingulate, and orbital cortex, and olfactory bulb), however, a few additional regions emerge (prelimbic cortex, amygdalar nucleus, pontine reticular nucleus) (**Supplement 1.2, Supplementary Figure 3, Supplementary Figure 4**).

### 2.4 Directly comparing the spatial pattern of SCT effects between species

We finally sought to achieve a direct test for the spatial convergence between SCT effects in humans and mice - separately for XXY and XYY. To achieve this test, we restricted analysis to a subset of brain regions with well-established homology between humans and mice based on both comparative structural and functional studies ^11,19–23^, and a model proposed by Swanson and colleagues ^21^ which maps neuroanatomical regions based on 6 cytoarchitectonic and MRI-derived human atlases and 3 cytoarchitectonic mouse atlases (as well as 2 rat atlases). Human brain atlases include Brodmann ^24,25^ and Swanson ^26^ (cytoarchitectonic), and Desikan-Killiany ^27^, Glasser ^11^, and the Allen Institute for Brain Science (MRI-based). The Allen Brain Institute ^28^, Hof ^29^, Paxinos and Franklin ^30^ were used as the mouse brain atlases. (**Table 1**) ^11,19–23^. For both the effect of an added X- or Y-chromosome, there was low global similarity between the effects on the human and mouse brain based on low correlation of effect sizes (added X cross-ROI r=−0.07; added Y cross-ROI r=−0.19; **Figure 5; Table 1**). However, the direction of effect sizes in each species highlighted several regions of convergent SCT effects on brain volume in both species (albeit with larger effect sizes in humans than mice). Specifically, XXY-induced cross-species volume reductions in the right amygdala, bilateral anterior cingulate and cerebellar cortex, left nucleus accumbens, right piriform cortex, right primary motor area, left retrosplenial area, left thalamus, and bilateral ventral orbital, and cross-species volume increases in the left posterior parietal association area, bilateral primary auditory and primary visual areas. XYY-induced crossspecies volume reductions in the bilateral amygdala, cerebellar cortex, left globus pallidus, left nucleus accumbens, bilateral piriform cortex, left primary motor area, and left temporal association area and cross-species volume increases in the right perirhinal area, left posterior association areas, left primary auditory and primary somatosensory areas (**Table 1**). The crossspecies similarities in SCT effects were also low for standardized beta values that were not TTV-corrected (**Supplement 1.3, Supplementary Figure 6, Supplementary Table 1**).

**Table 1.**
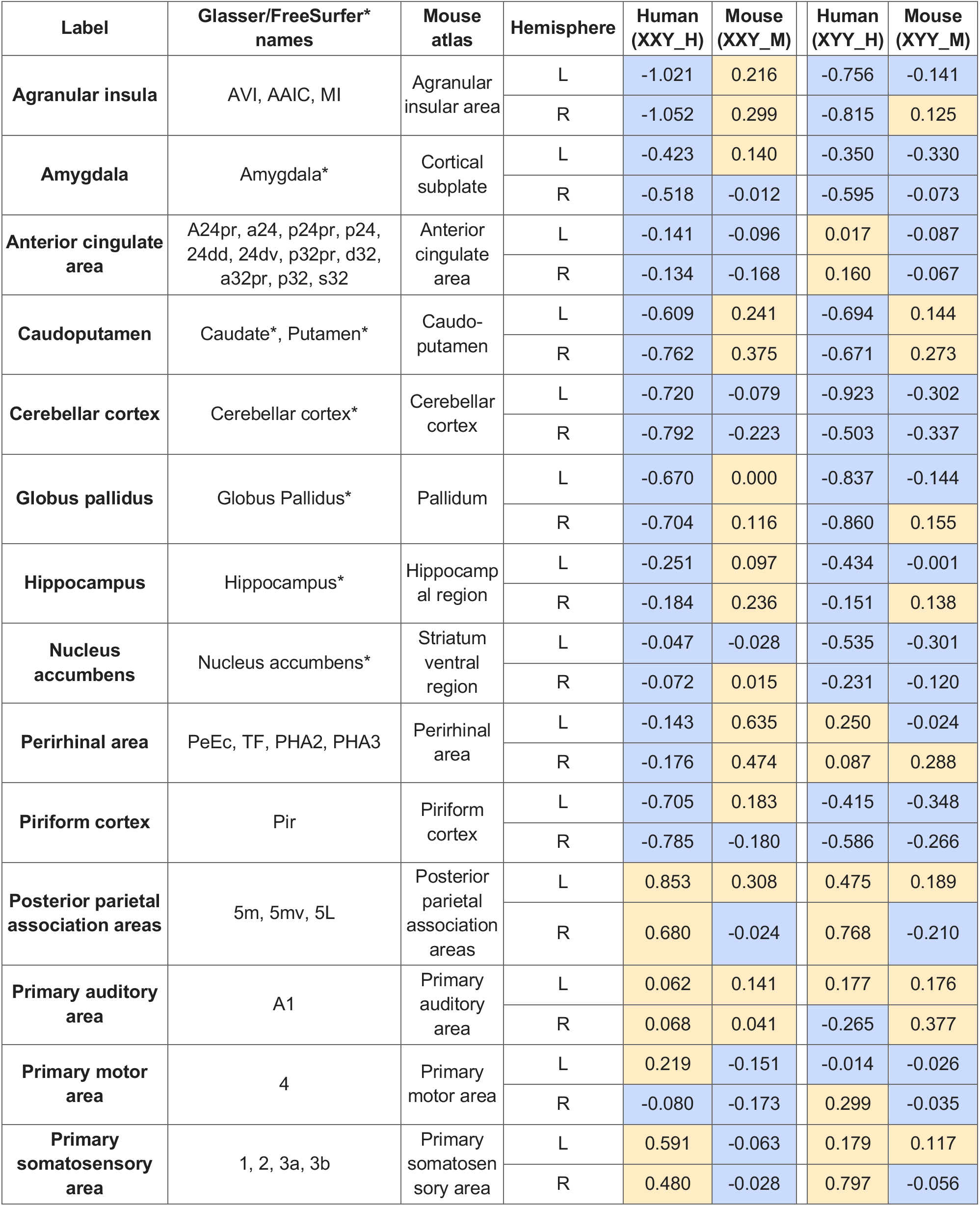
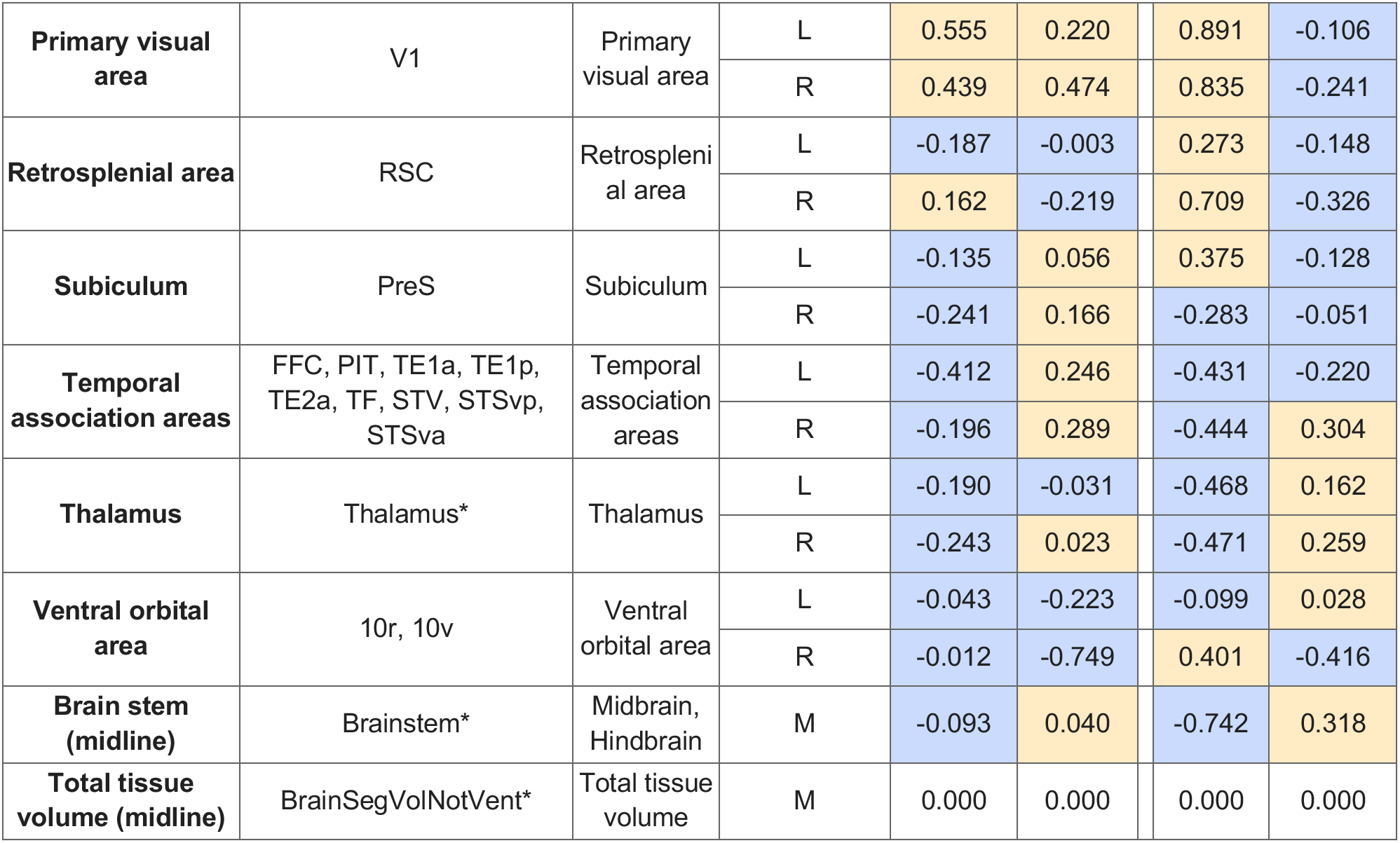
Mapping of homologous human-mouse brain regions (TTV-corrected standardized effect sizes). Blue shade highlights cells with negative standardized effect sizes, while peach highlights positive ones. Human = _Y; mouse = _M.

**Figure 5.**
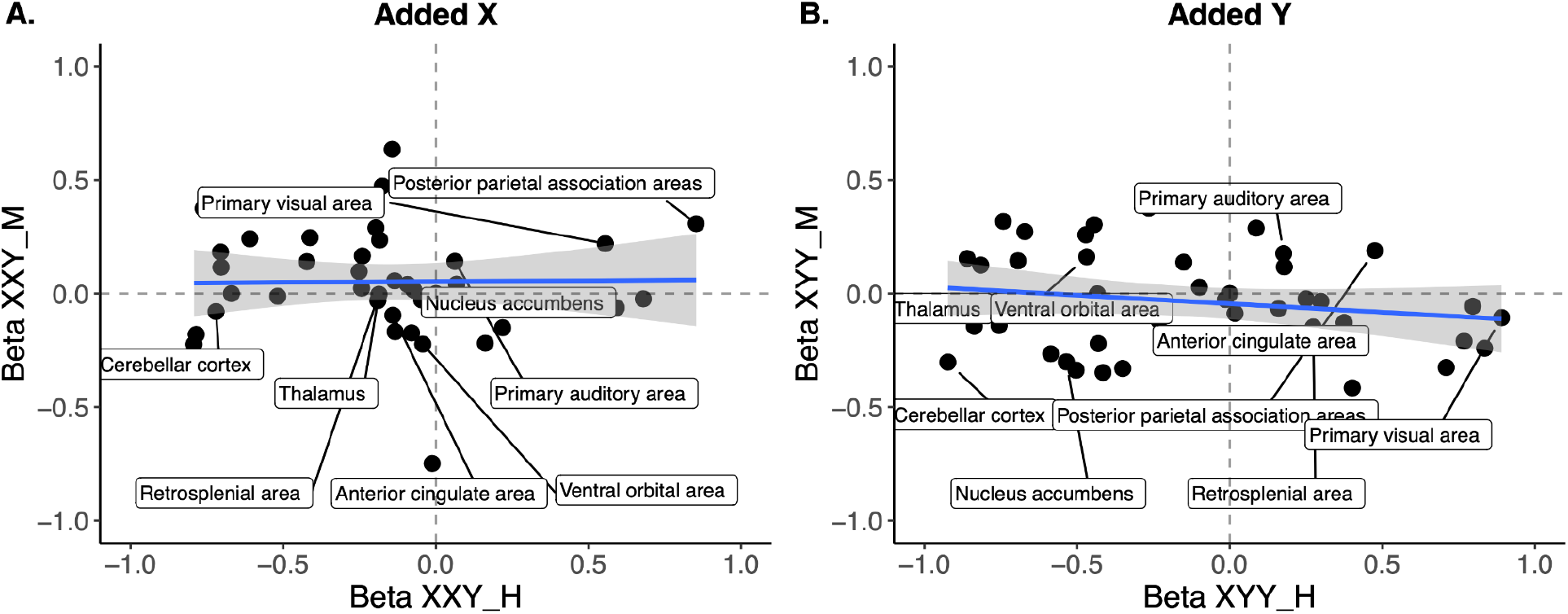
Correlation of effects of added X- or Y-chromosomes in human and mouse homologous brain regions. **A.** Standardized effect size correlation for the effect of added X-chromosome (cross-ROI r=−0.068) or B. added Y-chromosome (cross-ROI r=−0.183). Points are labeled only if they have the same directionality of standardized effect size in humans and mice (i.e., both positive or both negative). For simplicity, labels are for left hemisphere regions only as there were no large differences in laterality of effects. Human = _Y; mouse = _M.

## 3. Discussion

This study provides the first cross-species comparison of the effects of an added X- or Y-chromosome on mammalian brain anatomy, leveraging a unique sample of human youth with SCT and young adult transgenic SCT mice. Our findings substantially advance understanding of SCD effects on brain anatomy within and between species, and also take a first formalized step towards the broader topic of comparative neuroimaging of brain alterations between humans and mice. We consider each of these advances in turn below.

First, we detail the effects of XXY and XYY on global and regional brain volume in humans. We focus on the two most common SCTs, Klinefelter syndrome (XXY karyotype) and XYY syndrome (XYY karyotype)^31^ to model the effects of SCD on human brain anatomy. Our results align with those of two other SCA cohorts that have been studied to date ^8,32–36^ to provide an independent replication of several observations including (i) a decrease in TTV with carriage of a supernumerary X-chromosome in males, (ii) a trend towards greater TTV with carriage of a supernumerary Y-chromosome, and (iii) convergent effects of XXY and XYY on regional brain anatomy once these divergent global effects are controlled for ^8,32^. These convergent effects involve relative volumetric expansion of occipital and parietal cortices alongside relative volumetric contraction of insula and temporal cortices, as well as cerebellar, brainstem, and basal ganglia subregions. There is subtle variation within this general pattern of concordance between the effects of XXY and XYY on relative regional brain volume in humans (cross region correlation in effect size r>0.7). Although effects were present in both SCT groups, there were slightly more pronounced volume contraction of the putamen, amygdala, piriform, posterior cingulate, and frontal cortex in XXY vs XYY, and of the brainstem, auditory, and frontal cortex in XYY vs XXY. From a mechanistic perspective, the divergent effects of XXY and XYY on TTV could reflect those aspects of biology that distinguish these two SCTs. For example - XXY individuals have higher total chromatin content and gene count than XYY since the X-chromosome is larger and carries more genes than the Y-chromosome ^37^. Additionally, differences in the expressed number of Y-linked and X-linked genes (in particular those X-linked genes that escape X-inactivation) may contribute to observed differences in the effects of XXY and XYY ^38–41^. Altering the expression of either X- or Y-linked genes may affect regulation of transcription, translation, and protein stability ^41^. Secondary endocrine effects of XXY such as hypogonadism, low testosterone, and infertility ^42^, not seen in XYY, may also contribute to differences in brain anatomy between XXY and XYY. However, two observations suggest that the neuroanatomical changes seen in XXY are likely to reflect direct X-chromosome dosage rather than gonadal effects: individuals with XXX have similar neuroanatomical changes to those observed in XXY ^8^ and XO individuals have an inversion of these effects ^43,44^. Conversely, the shared neuroanatomical effects of XXY and XYY could reflect biological features that these two have in common. Specifically, XXY and XYY both lead to increased dosage of pseudoautosomal (PAR) genes, as well X-Y homolog genes, also referred to as gametologs. It is also theoretically possible that generic effects on chromosomal trisomy explain shared neuroanatomical changes in XXY and XYY, but this idea is countered by the fact that trisomy of chromosome 21 (Down Syndrome) induces regional brain volume changes that differ from those seen in XXY or XYY ^46,47^.

Second, we provide the first systematic analysis of X- and Y-chromosome dosage effects on regional brain anatomy in mice through the lens of SCT. We find that carriage of an extra X- or Y-chromosome is not associated with large effect size changes in either global or regional brain anatomy in mice, although some subtle effects are apparent. Of note, the subtle effects of XXY and XYY on regional brain volume in mice showed weak and moderate stability (respectively) between independent groups differing in gonadal background - suggesting that although subtle, some of these effects are indeed capturing direct impact of SCD on regional murine brain volume. Previous studies of SCT mice have found that carriage of an extra X-chromosome is associated with increased bodyweight and adiposity, while carriage of an extra Y-chromosome is associated with motor deficits ^4^. However, there are limited past studies of SCT effects on murine brain anatomy with which to compare our results, with the exception of a small series of studies in mice with X-monosomy and an XXY model that is derived in a different manner, and on a different strain background, relative to the SCT model studied here ^48,49^. Specifically, reduced retrosplenial, motor, and somatosensory cortex, main olfactory bulb volume, and increased anterior olfactory nucleus, lateral septum, bed nucleus of the stria terminalis, hypothalamus, and periaqueductal gray volume was observed in XXY compared to XY littermates ^48^. Interestingly, sensitivity to X-chromosome dose across XO, XX, XY, and XXY murine groups was observed in several brain regions including the periaqueductal gray, lateral septum, bed nucleus of the stria terminalis, amygdala, hypothalamus, and thalamus, whose volume increased with increasing X-dose, independent of the presence of the Y chromosome ^48,49^. We also observe enlarged periaqueductal gray volume in XXY relative to XY (and XYY relative to XY) mice, and reduced volume in the olfactory bulb, retrosplenial, and somatosensory cortex (at a relaxed threshold), similar to previous reports. Thus, our results combine with past work from two different murine models to suggest that volume of the periaqueductal gray, olfactory bulb, somatosensory and retrosplenial cortices in mice may be sensitive to X-chromosome count, where volume increases with X-dose. Potential drivers of X-chromosome dosage effect on the volume of these regions include PAR and non-PAR genes that escape X-inactivation, as well as interactions with gonadal hormones^1,9^.

Third, we formally compare SCT effects on brain anatomy between humans and mice at several complementary levels of analysis. We find a general lack of conservation for SCT effects between the two species. We observe globally divergent effects of XXY and XYY on human TTV, but minimal effects of either SCT group on TTV in mice. Regional volume alterations in human SCT are also not consistently replicated in murine SCT - either, in terms of their magnitude or anatomical localization. For example, dissimilarities between species were observed in the dorsal pallidum (decreased in humans and increased in mice) and the primary somatosensory cortex (increased in humans and decreased in mice). Humans and mice also differ in the within-species concordance of XXY and XYY effects, wherein the concordance of XXY and XYY effects is high in humans but low in mice. Overall, the magnitude of effects in human SCT are much larger and more spatially congruent between the X- and Y-chromosome than in mouse SCT. We speculate below on the potential proximal and distal mechanistic basis of discordance of SCT effects in the human and murine brain. Proximally, although sex chromosomes show privileged conservation between humans and mice ^3,50^, both in terms of chromosome size and specific gene expression ^51^, there are differences between species. The male specific region (MSY) of the Y-chromosome has a much larger proportion of ampliconic genes in the mouse compared to primates, driven by massive duplications of 3 gene families associated with testis-specific fertility ^52^. These changes have resulted in the degeneration of other genes, pushing the mouse MSY to diverge even more from the mouse X-chromosome ^52^ in gene content than the primate MSY has diverged from the primate X-chromosome. Although most of the expanded gene families on the mouse MSY are primarily expressed in the testes ^52^, they may indirectly explain some of the interspecies differences in Y-chromosome dosage on the brain. Although the X-chromosome has greater inter-species similarity than the Y, differences exist in the position of genes along the chromosome, and in the presence of certain newly acquired ampliconic genes ^39,53^ which may contribute to differences in SCT effects across species. Mice also differ from humans in X-Y gene homology. Mice only have 1 small PAR, which allows for recombination between sex chromosomes, while humans have two larger regions containing more genes than the mouse PAR ^51^. Numerous PAR genes in humans are expressed to a higher degree in males than females ^54^. Furthermore, relative to other placental mammals, including humans, mice have lower expression levels of Y gametolog genes ^55^. Finally, the process of X-chromosome inactivation (XCI) differs between species, as does the degree of escape from X inactivation. XCI initiates early in murine development (at the 2-cell stage), wherein *Xist*, the transcription factor responsible for initiating XCI, coats the paternal X-chromosome ^39^. The paternal X-chromosome is then reactivated at the mid-blastocyst stage, where XCI is reinstated and random. Conversely, in humans there is no paternal imprinting of XCI, and random silencing only occurs later in development (50 cell stage) ^39,51^. Humans and mice also differ in escape from XCI, wherein 12-15% of human and 3-5% of mouse genes escape consistently, while an additional 8-10% of human genes and 4% of mouse genes have variable tissue specific escape ^37,39,56^. Given the putative importance of escape genes on brain volume ^57^, these may play a key role in mediating species differences in X-chromosome dosage effects on the brain. Distally, there are also differences between humans and mice in the autosomal context that the sex chromosomes are operating within, and the evolutionary, developmental, and environmental contexts within which these genetic effects unfold.

Despite the general lack of similarity between SCT effects on the murine and human brain, our study design did identify a handful of regions that show convergent directional effects of SCT between XXY and XYY and between species. These include decreased cerebellum, cingulate, and orbital cortex volume and increased parietal association cortex volume. These regions represent high priorities for follow-up studies aimed at understanding mechanistic effects of SCT on the brain, improving the translation between species. Deeper investigation of the regions in which effects had opposite directionality between species may help us understand species differences in the effects of SCD on brain anatomy.

Finally, our attempt to formalize a test of convergent anatomical effects of SCT on the human and murine brain provides the first worked example of a more broadly valuable study design whereby common neuroimaging methods provide a highly structured and standardized framework for comparing brain changes between species. There have been huge advances in computational morphometry of the brain within humans ^11,58^ and mice ^9^, which have been used to tackle similar questions in each species (e.g., mapping genetic effects) - but not yet in a combined fashion as attempted in the current study. The effort undertaken here raises several important insights for future comparative neuroimaging. We employed a parcellation-based approach, looking for similarities and differences in the directionality of volume changes due to SCT. However, there are different ways of defining the units of analysis within each species and different ways of mapping between them that go beyond classical anatomical nomenclature (which is subject to change based on the parcellation used ^59^). Defining homology based on dominant functional systems - i.e., visual system in humans to the olfactory system in mice - goes beyond region-to-region comparison and considers an animal’s environmental and evolutionary niche ^60^. This may be an intriguing avenue of future research as we do observe effects of SCT in the human visual system and the mouse olfactory system. Alignment across species may also be achieved by comparing patterns of homologous gene expression ^61^ which reflect distinct patterns of neuroanatomical organization both at the molecular and morphometric level ^62–64^. Furthermore, gene expression patterns are generally conserved between species ^65^, particularly in regions that are comparable between species, such as the motor cortex, striatum, and cerebellum ^66^. Although global patterns of gene expression are comparable, single-cell transcriptional analyses have revealed important species divergence in non-neuronal (glial cell, and neurotransmitter system) expression patterns between humans and mice ^65^. Alignment may also be achieved by comparing connectivity patterns between brain regions using functional or diffusion MRI, which are less time consuming than the gold-standard tracer studies ^67^. Previous work has identified both homologous and divergent connections when comparing human and macaque cortical circuits ^68–70^, as well as human, macaque, and mouse cortico-striatal circuits ^23^. Considering differences in the developmental timing of the brain across species may provide an alternative lens through which to establish homology. Additionally, multiple outputs can be used as tests for agreement between species including (i) similarities at the global and local level, (ii) the magnitude and direction of the effects, (iii) effects at a region-by-region level or across multiple regions. Furthermore, defining “large” vs. “small” effects within each species may be important for interpreting the significance of observed differences. While structural MRI brings new opportunities to formalize cross-species testing as compared to other modalities such as behavior, there are inevitable species differences in lifestyle and sources of variance between laboratory mice (both inbred and outbred ^71^) and outbred humans.

Our results should be considered alongside some limitations. There may be ascertainment bias in the sample of human youths with SCT recruited in this study, as patients presenting with more severe phenotypes are more likely to seek clinical attention and enroll in research ^72,73^. Additionally, there may be confounding effects due environmental exposure in humans that would not be present in mice as they are reared in a tightly controlled laboratory environment. Furthermore, the influence of endocrine factors or sex steroids may further confound effects in humans. For example, individuals with XXY typically have low testosterone levels, which can be corrected via hormone therapy ^74^, both of which may impact brain anatomy ^75^. We base our species comparison solely on structure volume, relying on potentially inconsistent nomenclature of areas across species. Although the brain of both species is similarly organized into cortical, subcortical, cerebellar, and brainstem nuclei, obtaining homology at a finer grain level is challenging, particularly for multimodal association cortices which have diverged between species, more than other regions ^23,76^. There is great variability in the size and organization of the neocortex across mammals, which likely points to a convergence of evolutionary pressures on this structure. Understanding interspecies differences in the cortex, although challenging, may aid in our understanding of evolution. As discussed above, novel methods with which to formalize translation via the use of transcriptional similarity ^61^, or connectivity ^23,77^ have shown great promise, and should be applied to the study of SCD in future work. Furthermore, integration of these findings across other mammals could enhance our understanding of the evolutionary pressures driving species differences in SCD effects on the brain; however, generating animal models (other than the mouse) with additional sex chromosomes may be a challenging endeavor.

In conclusion, our study shows that the effects of added X- or Y-chromosomes in humans have diverging effects on total tissue volume, but large, and spatially similar effects on regional brain volume. In contrast, we observe subtle, and spatially dissimilar effects due to added X- or Y-chromosomes in the murine brain. However, our screen of regional brain anatomy does identify several brain regions which show similar directionality of volume change due to SCT in both humans and mice (e.g., cerebellum, parietal, cingulate, orbital cortex). These regions represent high-priority targets for future translational dissection of SCD effects on mammalian brain development, which may allow for a deeper mechanistic understanding of how SCD affects neuroanatomy. Establishing methods for cross-species comparison provides a promising avenue for better understanding the strengths and limitations of animal models in the study of the human brain in health and disease.

## Supporting information

Supplementary results, figures, and tables

## Acknowledgements

This study was supported by the intramural research program of the National Institute of Mental Health (project funding: 1ZIAMH002949-03; clinical trials identifier: NCT00001246; clinicaltrials.gov; protocol: 89-M-0006) and the National Institute of Child Health and Disease (R01HD100298). EG also receives salary support from the Fonds de Recherche du Québec en Santé. We would like to acknowledge Alyssa Warling and Ethan Whitman for their contributions to the collection of the human XXY and XYY data.

## Author contributions

Conceptualization, writing - original draft, EG, AR, JPL;; supervision, AR, JPL; project administration, EG, AR, JPL, AMG, XY, APA; methodology, software, and formal analysis, EG, AB, SL, EL, AR, JPL; visualization, EG, AB; human data collection, LC, ET, JB, FL; mouse data collection, LQ, HH; funding acquisition, AR, JPL, AMG, XY, APA; writing - review & editing, all authors.

## Inclusion and diversity

One or more of the authors of this paper self-identifies as an underrepresented ethnic minority in science. While citing references scientifically relevant to this work, we also tried to promote gender balance in our reference list, although unfortunately we did not achieve it.

## Resource Availability

### Lead contact

Further information and requests for resources should be directed to and will be fulfilled by the lead contacts, Dr. Armin Raznahan and Dr. Jason Lerch.

### Materials availability

This study did not generate new unique reagents.

### Data and code availability

All raw data reported in the paper are available upon request to the corresponding authors. Input regional volume measures for humans and mice, as well as original code for statistical analyses have been deposited to github and are publicly available here: https://github.com/elisaguma/SCT-human-mouse-volumes/tree/main/.

## Key resources table

| REAGENT or RESOURCE | SOURCE | IDENTIFIER |
| --- | --- | --- |
| **Experimental models: Organisms/strains** |  |  |
| Mouse: MF1 Sex Chromosome Trisomy (SCT) mouse model | Bred at UCLA (Art Arnold Lab) | https://arnoldlab.ibp.ucla.edu/models/ |
| **Software and algorithms** |  |  |
| R version 3.6.0. | https://cran.r-project.org/bin/windows/base/old/3.6.0/ | RRID:SCR_001905 |
| RMINC library | https://github.com/Mouse-Imaging-Centre/RMINC | N/A |
| Data.tree library | https://cran.r-project.org/web/packages/data.tree/vignettes/data.tree.html | N/A |

## 4. STAR Methods

### 4.1 EXPERIMENTAL MODEL AND SUBJECT DETAILS

#### 4.1.1 Participants

The human sample in this study includes youth (aged 6-25 years) with one of two sex chromosome aneuploidies (XXY_H or XYY_H), each with their own completely independent euploidic control (XY_H) sample (XXY_H, n=99, XY_H control, n=82, and XYY_H, n=34, XY_H controls, n=37). Independent control groups were acquired due to a scanner upgrade occurring between bouts of data collection. Additionally, the independence of control groups reduces the risk that effects could be driven by the controls (rather than the aneuploidies). Participants from the SCA groups were recruited through the National Institute of Health (NIH) website and parental support groups, while healthy participants were recruited from the NIH Healthy Volunteer office. All participants underwent structural magnetic resonance imaging (MRI), and had normal radiological reports, with no history of brain injury or neurological disorders. Supernumerary X- and Y-chromosome carriage was confirmed by karyotype, and non-mosaicism was confirmed through visualization of 50 metaphase spreads in peripheral blood for XXY_H and XYY_H participants. XY_H controls were screened to exclude a history of psychiatric or neurodevelopmental disorders. This study was approved by the NIH Combined Neuroscience Institutional Review Board. All participants gave consent or assent, as appropriate, and all protocols were completed at the NIH Clinical Center in Bethesda, Maryland (see **Table 2** for demographics information).

**Table 2.** Demographics for human sample.

| Characteristic | XY | XXY | Statistics | XY | XYY | Statistics |
| --- | --- | --- | --- | --- | --- | --- |
| Sample size | 82 | 99 |  | 37 | 34 |  |
| Age (years) |  |  |  |  |  |  |
| Mean | 16.41 | 16.38 | F(1,179)=0.001, p=0.973 | 14.74 | 15.49 | F(1,69)=0.414, p=0.522 |
| SD | 5.77 | 4.78 |  | 4.53 | 5.33 |  |
| Range | 6.03 - 24.99 | 6.67 - 25.8 |  | 5.6 - 24.7 | 7 - 25.9 |  |
| Full Scale IQ |  |  |  |  |  |  |
| Mean | 114.9 | 93.25 |  | 117.62 | 87.15 |  |
| SD | 12.59 | 12.22 | F(1,179)=137,<br>p<2e-16 ** | 9.4 | 12.40a | F(1,69)=137.6,<br>p<2e-16 ** |
| Range | 83-146 | 64-127 |  | 97-136 | 62-111 |  |
| Socioeconomic status (SES) |  |  |  |  |  |  |
| Mean | 38.39 | 47.34 | F(1,179)=9.54<br>3 0.0023 * | 38.89 | 54.91 | F(1,69)=17.16,<br>p=9.57e-05 ** |
| SD | 17.87 | 20.6 |  | 14.47 | 18.04 |  |
| Range | 20-95 | 20-120 |  | 20-70 | 20-95 |  |

\*P < 0.01 \*\*P < 0.001 for anova test of significant difference between groups (XY vs XXY or XY vs XYY)

#### 4.1.2 Mice

The Sex Chromosome Trisomy (SCT) mouse model ^4^ is generated by mating XXY^−^ gonadal females with *XY*^−^(*Sry*^+^) gonadal males. The testis-determining gene, *Sry*, is deleted from the Y^−^ chromosome ^78^, preventing formation of testes and causing development of ovaries in XXY^−^ mothers. The *XY*^−^(*Sry*^+^) father possesses an *Sry* transgene on chromosome 3, causing differentiation of testes. ^79–81^. The XXY^−^ mothers produce X and XY^−^ eggs, while the XY^−^(*Sry*^+^) fathers produce X or Y^−^ sperm cells each with or without *Sry*^4^. This yields eight potential genotypes: XX, XY, XXY, and XYY, each genotype with either testes or ovaries. In order to mirror the human SCT groups, gonadal male XY_M (n=18), XXY_M (n=17), and XYY_M (n=20) mice were included in this study. All experiments were approved by the UCLA Animal Research Committee. Mice were group-housed at the UCLA animal facility based on gonadal sex, maintained at 23°C on a 12:12 light/dark cycle, and fed regular chow diet with 5% fat *ad libitum* (LabDiet 5001, St. Louis, MO, USA)

### 4.2 METHODS DETAIL

#### 4.2.1 Human Neuroimaging

##### Neuroimaging data acquisition

High-resolution MPRAGE T1-weighted (T1w) sMRI scans was acquired on each participant using a MR750 3-Tesla (General Electric) whole-body scanner with a 32-channel head coil (176 contiguous sagittal slices with 256 × 256 in-plane matrix and 1 mm slice thickness yielding 1 mm isotropic voxels). T1w sMRI scans were converted from DICOM to Nifti and organized according to the Brain Imaging Data Structure (BIDS) using heudiconv.

##### Data processing

###### Cortical segmentation

The BIDS compatible Freesurfer pipeline (version 7.1.0) ^12^ was used for cortical and subcortical segmentation of the T1w MRI scans (https://github.com/Shotgunosine/freesurfer/tree/fs_7), which is documented and freely available for download online (http://surfer.nmr.mgh.harvard.edu/). The technical details of these procedures are described in prior publications ^1–14^. Cortical surface reconstruction was achieved using the automated *recon-all* processing stream. From the surfaces, a number of features were extracted *mri_anatomical_stats* utility (cortical thickness, surface area, cortical volume, mean curvature, gaussian curvature, intrinsic curvature index, and folding index). In order to use a homologous measure across the human and mouse analyses, we focused on cortical volume. Vertex-level cortical volume measures were averaged across 360 cortical parcels from the Glasser Human Connectome Project regional parcellation, a multimodally-informed atlas ^11^. The Euler number was extracted from each individual’s cortical reconstruction. This is thought to reflect image quality and topological complexity. Participants with a Euler number less than −217 were excluded from statistical analyses as this has been previously shown to be a robust quality control threshold and a proxy for in-scanner motion ^82^.

###### Subcortical segmentation

In automatic subcortical segmentation, also part of the automated *recon-all* processing stream, each voxel in the normalized brain volume is assigned one of about 40 labels using Freesurfer “aseg” feature (version 7.1.0; see ^83,84^ for full details). Briefly, all T1w scans were affinely registered to MNI space followed by initial volumetric labeling and B1 bias field correction. Next, images T1w images were nonlinearly registered to MNI space followed by a final labeling of the volume. Freesurfer uses probabilistic estimations based on Markov random fields ^85^ to automatically assign a label to each T1w image voxel based on a training set (previously manually delineated brains). The boundaries of regions are estimated based on expected shapes of structural and signal intensity from the T1w image. Each scan was visually inspected to exclude images that were corrupted by motion artifacts or had segmentation errors.

#### 4.2.2 Mouse Neuroimaging

##### Data acquisition

Adult mice (postnatal day 65) were anesthetized, transcardially perfused, and decapitated at UCLA as previously described ^86^, and shipped for scanning at the University of Toronto. Brains (kept within the skulls) were placed in a solution of paraformaldehyde (PFA) and Prohance^®^ (gadoteridol, Bracco Diagnostics Inc., Princeton, NJ) for fixation. A multichannel 7.0-T scanner with a 40-cm diameter bore magnet (Varian Inc., Palo Alto, CA) was used to acquire MR images at 40-μm-isotropic resolution ^87^ for 16 brain samples concurrently ^88^. Scan parameters: T2W 3D FSE cylindrical *k*-space acquisition sequence, TR/TE/ETL = 350 ms/12 ms/6, two averages, FOV/matrix-size = 20 × 20 × 25 mm/ 504 × 504 × 630, total-imaging-time = 14 h ^87^.

##### Data processing

Structural MRI images (n=55) were aligned by unbiased deformation based morphometry using a previously described ^89,90^ registration pipeline ^91–93^. Briefly, all individual scans are registered together using affine and nonlinear registration to create a study specific average/template, from which log-transformed Jacobian determinants can be calculated ^94^, quantifying the voxel-wise volumetric differences between each individual image and the study specific average/template. The MAGeT brain algorithm ^95,96^ was used to segment images according to a published atlas of 355 unique regions ^13–16^.

### 4.3 QUANTIFICATION AND STATISTICAL ANALYSIS

#### 4.3.1 Effects of sex chromosome aneuploidy on regional and voxel-wise brain volume

##### Human

All statistical analyses were performed in R version 3.6.0. Volumes for each aneuploidy group were z-scored using the mean and standard deviation measures of their independent XY_H control group. For each case-control pair, we evaluated the effects of SCT on the volume of each cortical and subcortical brain structure using a linear model to assess the effect of group (β1: XXY_H vs XY_H or XYY_H vs XY_H), with mean-centered age (β2), and TTV (β3) as covariates (ε: error term). Test statistics were corrected for multiple comparisons using the False Discovery Rate (FDR) correction ^17,18^ with *q* (the expected proportion of false positives) set at 0.05. The formula, using ROI_volume as the example region:

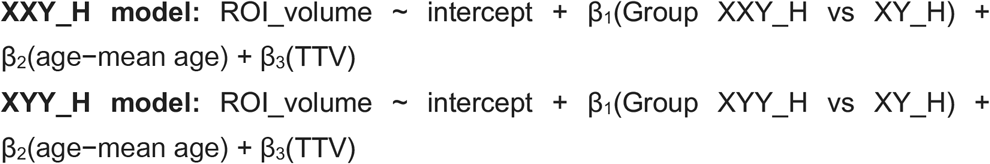

##### Mouse

To mirror the human analyses, volumes for all groups were z-scored using the mean and standard deviation of the XY_M control group. Effects of SCD on brain volume were again assessed using a linear model testing for the effect of group (β1: XXY_M or XYY_M vs XY_M control), with total brain volume (β2) as a covariate (ε: error term). Since there was a single control group, we ran a single linear model, comparing each SCT group to the XY control (set as the reference group). We did not covary for age in this model as the mice were of the same age.

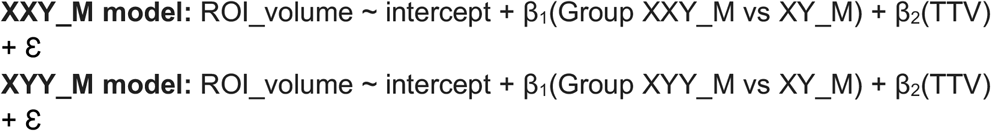

We performed a supplemental analysis to test for the reproducibility of effects in XY, XXY, and XYY mice with ovaries (XY [n=24], XXY [n=19], and XYY [n=22]). We computed the standardized effect size (β_1_) for the effect of an added X- or Y-chromosome and correlated them with the standardized effect sizes computed for mice with testes across all brain regions. Additionally, in both species, analyses were repeated without TTV correction (**Supplement 1.1**).

#### 4.3.2 Spatial correlation and convergence of effects between XXY and XYY groups

##### Human

To assess the spatial convergence of the effects of adding an X or adding a Y chromosome on brain volume, the β_1_ value from the XXY_H and XYY_H models was correlated across all brain regions. Regions that were either consistently increased or decreased due to the addition of an X or Y chromosome were identified based on the sign of the β1 coefficients in statistically significant regions.

##### Mouse

Since each mouse aneuploidy group did not have their own independent control, we performed a bootstrap resampling without replacement to generate two independent control groups as follows: we repeatedly split (n=1000) the XY control group (n=18) in half, sample A (n=9) and sample B (n=9), without replacement. Sample A was used to calculate z-scored brain volumes for the XXY mice, while sample B was used for the XYY mice. For each control-split, and across all brain regions, a linear model to assess for group effects, with correction for total brain volume. For each control split, the β1 was stored.

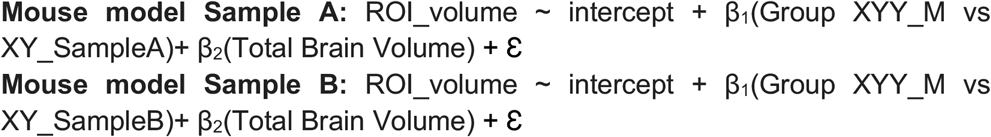

The β_1_ value of each brain region for sample A was correlated to that of Sample B across every region in the brain, yielding a 1000 × 1000 correlation matrix for each of the control splits. Next, we identified which brain regions had either a consistently positive or a consistently negative β1 value across ≥95% of the splits (950/1000 splits) for both the XXY_M (with XY_M Sample A) and XYY_M (with XY_M Sample B) groups. In both species, analyses were repeated without TTV correction (**Supplement 1.2**).

#### 4.3.3 Cross-species comparison for the effects of an added X- or Y-chromosome

We identified 21 brain regions determined to be homologous between humans and mice based on a framework presented by Swanson and Hoff ^21^, in which human, rat, and mouse brain regions are compared across a number of different anatomical and histological atlases, as well as the work of others ^19,20,61^ (**Table 1**). We repeated the procedure described in **4.3.1** to generate β-values for the effect of group (covarying for total tissue volume for both species and mean-centered age for humans) using z-scored volumes relative to the XY_H and XY_M controls. The β-values for the effect of adding an X-chromosome were correlated across species (XXY_H with XXY_M), as were those for adding a Y-chromosome (XYY_H with XYY_M) (**Figure 4**). In both species, analyses were repeated without TTV correction (**Supplement 1.3**).

## Legends for any Supplemental Excel Tables or files

**Supplementary Figure 1. Effect of added X- or Y-chromosomes on regional brain volume in humans and mice, without total tissue volume (TTV) correction.** Distribution of beta values for the effect of aneuploidy are displayed for humans (**A**) and for mice (**B**). **C.** Representative views for the cortical Glasser atlas for lateral and medial slices of the left and right hemispheres, and subcortical FreeSurfer atlas with both axial and sagittal views. Unthresholded (left) and significant (q<0.05; right) beta values for the effect of added X (**D**) and added Y (**E**) in the human brain. **F.** Mouse brain coronal slices with the representative atlas on the left, followed by untresholded, then thresholded (p<0.05) beta values for the effect of added X, followed by untresholded, then thresholded (p<0.05) effects of added Y.

***Supplementary Figure 2. Similarity of SCT effects on neuroanatomy across gonadal background.** Correlation of standardized effect sizes for the effect of an added (**A**) X chromosome (XXY vs XY)* (r=0.22) *and added (**B**) Y chromosome (XXY vs XY)* (r=0.56) *for mice with testicular background or ovarian background*.

**Supplementary Figure 3. Spatial convergence between added X- or Y-chromosomes on the human and mouse brain without total tissue volume (TTV) correction**. **A**. Human brain regions whose volume was either increased (red) or decreased (blue) in both aneuploidy groups. **B**. Correlation between beta values for the XXY_H and XYY_H group across all brain regions. **C**. Mouse brain regions whose volume was consistently increased (red colours) or decreased (blue colours) in 95% (950/1000 splits) of the XY_M control splits in both aneuploidy groups based on bootstrap analysis. **D**. Correlation between beta values for a representative split (34) for XXY_M and XYY_M groups across all brain regions. All plots use the beta values for the effect of aneuploidy with total brain volume correction.

**Supplementary Figure 4. Bootstrap analysis shows low spatial correlation for effects of an added X- or Y-chromosome in the mouse brain.** The coefficient for the correlation between cross-ROI standardized effect size (β) for XXY_M (relative to XY_M sample A) and XYY_M (relative to XY_M sample B) both with TTV correction (**A**) and without (**B**).

**Supplementary Figure 5. Convergently increased or decreased regions in humans and mice without total tissue volume (TTV) correction.** Venn diagram of overlapping ROIs highlighting the regions congruently (for both XXY and XYY) increased or decreased in humans and mice. Brain regions that were statistically significantly impacted by both XXY and XYY in humans alone (<5%FDR) are listed in the red (increased volume) and light blue (decreased volume) boxes on the left. Brain regions that showed significantly convergent impacts of XXY and XYY (based on bootstrap resampling) in mice alone are listed in the yellow (increased volume) and dark blue (decreased volume) boxes. There were no regions showing convergence (increased or decreased), or opposite (increased in one species and decreased in another, or vice versa) highlighted by the “NA” in the intersecting cells.

**Supplementary Figure 6. Correlation of effects of added X- or Y-chromosomes in human and mouse homologous brain regions when total tissue volume (TTV) is not controlled for. A.** Standardized effect size correlation for the effect of added X-chromosome (cross-ROI r=−0.06) or **B**. added Y-chromosome (cross-ROI r=-0.19). Points are labeled only if they have the same directionality of standardized effect size in humans and mice (i.e., both positive or both negative). For simplicity, labels are provided only for left hemisphere regions given that effects were similar across hemispheres.

**Supplementary Table 1. Mapping of homologous human-mouse brain regions (standardized effect sizes with TTV correction).** Blue shade highlights cells with negative standardized effect sizes, while peach highlights positive ones.

**Supplementary Table 2. Summary table for results for human XXY and XYY results with TTV correction for each ROI.** Annotations are made to highlight significance, and congruence in directionality across groups. (for attached document: H_Lm_wTBV_outputs_summary_table.xlsx)

**Supplementary Table 3. Summary table for results for human XXY and XYY results without TTV correction for each ROI.** (for attached document: H_Lm_noTBV_outputs_summary_table.xlsx)

**Supplementary Table 4. Summary table for results for mouse XXY and XYY results with TTV correction for each ROI.** (for attached document: M_Lm_wTBV_outputs_summary_table.xlsx)

**Supplementary Table 5. Summary table for results for mouse XXY and XYY results without TTV correction for each ROI.** (for attached document: M_Lm_noTBV_outputs_summary_table.xlsx)

