## Supplementary results, figures, and tables for "A cross-species study of sex chromosome dosage effects on mammalian brain anatomy"

### Supplementary Materials

#### 1. Results

##### 1.1 Regional brain volume changes due to added X- or Y- chromosome in humans and mice without total tissue volume (TTV) correction

In humans, without TTV correction, the effect of an added X-chromosome on standardized effects sizes across all brain regions were largely negative, (range of $\beta$=-1.448 to 0.464), while those for an added Y-chromosome were more evenly distributed, with both positive and negative effects (range of $\beta$=-1.239 to 1.564) (**Supplementary Figure 1A)**. Indeed, mean standardized effect sizes for XXY_H ($\beta$=-0.422) were significantly smaller than those for the XYY_H group ($\beta$=0.184), which were close to zero (comparison of effect size distributions, t=-20.838, df=714.58, p< 2.2e-16). The range of standardized effect sizes was also significantly different (Levene's test, F(1, 754)=18.145, p=2.306e-05). In contrast, in mice, the range of standardized effect sizes across brain regions was similar for the effect of an added X- ($\beta$=-0.848 to 0.385) and added Y-chromosome ($\beta$=-0.792 to 0.313) (**Supplementary Figure 1B)**. In contrast to the TTV corrected effects, the range and mean standardized effect sizes for XXY_M ($\beta$=-0.186) and XYY_M ($\beta$=-0.228) were shifted towards more negative, and significantly different from zero (t=2.799, df=904.82, p=0.005), but not significantly different from each other (Levene's test: F(1,906)=6e$^{-4}$, p=0.981).

Without TTV correction, large decreases in regional brain volume were observed in XXY_H driven by global volume reductions identified in **2.1**. These include widespread reductions in insula, orbitofrontal, fusiform face, frontal, piriform, visual, temporal cortex, cingulate, somatosensory association cortices, and in the cerebellum cortex, putamen, pallidum, amygdala, brainstem, thalamus, hippocampus, and caudate. Volume increases were observed in the somatosensory cortex (5m), visual cortex, paracentral lobule, and posterior cingulate cortex (POS2). Interestingly, the spatial patterning and directionality of XYY_H effects on regional brain anatomy were similar to results with TTV correction applied (**2.2**). Volume of the insula, fusiform face complex, inferior frontal cortex, and right pallidum was decreased, while volume of the visual, parieto-occipital, posterior cingulate, retrosplenial cortex, and somatosensory association cortex was increased. The effects of added X- or Y-chromosomes were not as spatially similar without TTV correction due to a global volume reduction in XXY_H.

In mice, non-significant (unthresholded) brain-wide reductions were observed in both XXY_M and XYY_M, with increased volume in the striatum, auditory and visual cortices, and brainstem of XXY_M, and increased mid- and hind-brain volume in the XYY_M. At an uncorrected (t=2.042, p<0.05) threshold, we observed volume decreases in the left main olfactory bulb granular layer ($\beta$=-0.662) and accessory olfactory bulb glomerular layers ($\beta$=-0.790), as well as the right cingulate cortex (area 29c) ($\beta$=-0.780), ventral orbital area ($\beta$=-0.871), and right primary somatosensory cortex ($\beta$=-0.643) for XXY_M. In XYY_M, volume decreases were observed in the left piriform-amygdalar area ($\beta$=-0.600), right primary somatosensory area (also decreased in XXY_M; $\beta$=-0.726), right cuneate nucleus ($\beta$=-0.675), and left paramedian lobule of the cerebellum ($\beta$=-0.851). Results for XXY_M and XYY_M without TTV were similar to observations made with TTV correction (**Supplementary Figure 1F**).

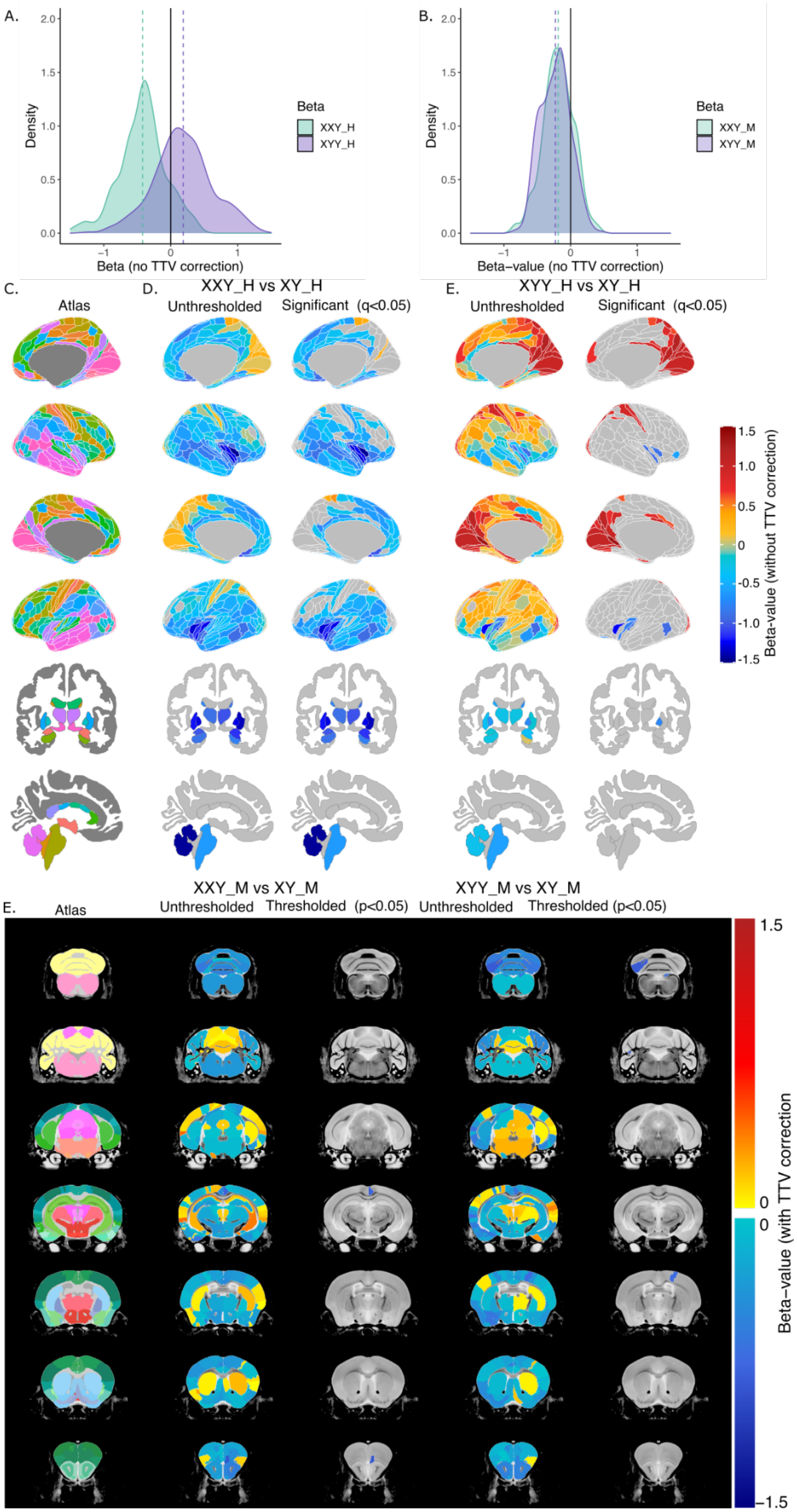

A.
Density
Beta (no TTV correction)
Beta
XXY_H
XYY_H
B.
Beta–value (no TTV correction)
XXY_M
XYY_M
C.
Atlas
D.
XXY_H vs XY_H
Unthresholded
Significant (q<0.05)
E.
XYY_H vs XY_H
Beta-value (without TTV correction)
XXY_M vs XY_M
Thresholded (p<0.05)
XYY_M vs XY_M
Beta–value (with TTV correction

***Supplementary Figure 1. Effect of added X- or Y- chromosomes on regional brain volume in humans and mice, without total tissue volume (TTV) correction.*** *Distribution of beta values for the effect of aneuploidy are displayed for humans (**A**) and for mice (**B**). **C.** Representative views for the cortical Glasser atlas for lateral and medial slices of the left and right hemispheres, and subcortical FreeSurfer atlas with both axial and sagittal views. Unthresholded (left) and significant (q<0.05; right) beta values for the effect of added X (**D**) and added Y (**E**) in the human brain. **F.** Mouse brain coronal slices with the representative atlas on the left, followed by untresholded, then thresholded (p<0.05) beta values for the effect of added X, followed by untresholded, then thresholded (p<0.05) effects of added Y.*

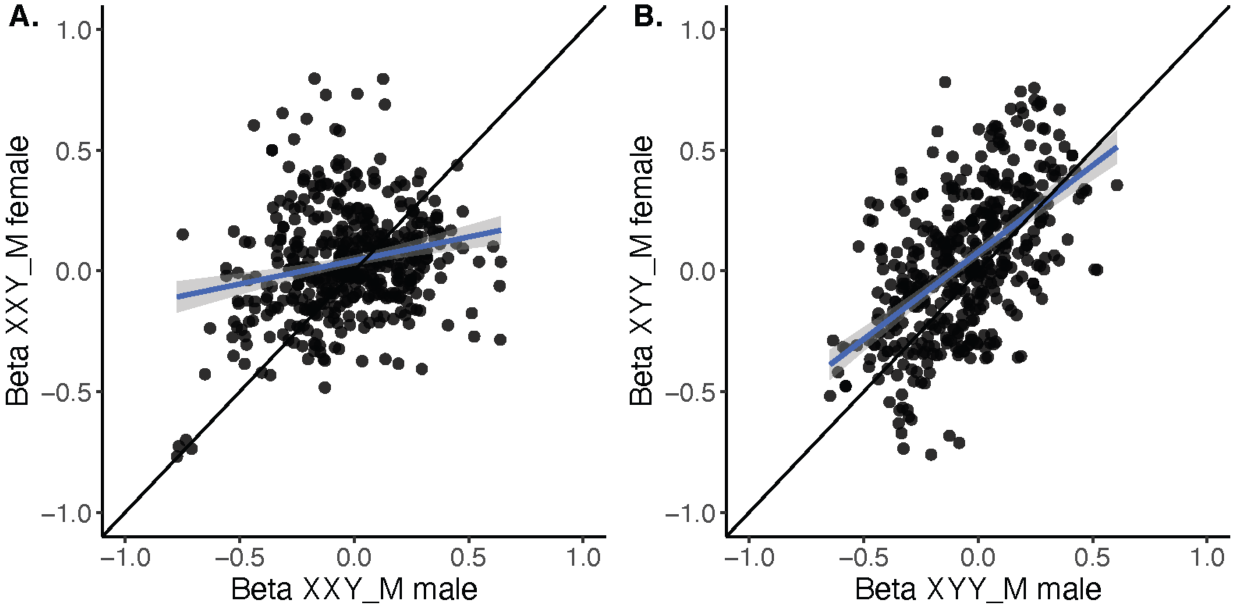


***Supplementary Figure 2. Similarity of SCT effects on neuroanatomy across gonadal background.*** *Correlation of standardized effect sizes for the effect of an added (**A**) X chromosome (XXY vs XY)* (r=0.22) *and added (**B**) Y chromosome (XXY vs XY)* (r=0.56) *for mice with testicular background or ovarian background*.

**1.2 Spatial similarity of added X- or Y-chromosome effects without TTV correction in humans but not mice**

A similar pattern of altered brain anatomy was observed in both human SCA groups, although to a lesser degree than in the TTV corrected analysis (**2.3**). A few subregions within the parietal and visual cortices (n=6 regions) were significantly increased in both XXY_H and XYY_H, while the pallidum, insula, temporal, and inferior frontal (inferior frontal gyrus and fusiform face complex) were both decreased due to SCA (n=11 regions) (**Supplementary Figure 3A**). As with the TTV corrected analysis, the overall similarity of effects was still high (cross-ROI r=0.73) in humans (**Supplementary Figure 3B**). In the mouse, the bootstrap analysis revealed no brain regions consistently increased (>95% of the time) in both SCA groups (**Supplementary Figure 3C)**. However, several regions were consistently and congruently decreased across both murine SCT groups, including the right cingulate cortex (29c), right primary somatosensory cortex, ventrolateral orbital areas, paramedian lobule of the cerebellum, and several olfactory bulb regions, which were also consistently and congruently decreased in the TTV corrected analyses. However, there were congruent decreases in the prelimbic area, left medial amygdalar nucleus, left crus 2, and right pontine reticular nucleus of both XXY and XYY mice, which were not observed when correcting for TTV effects. The distribution of cross-ROI correlations was centered close to zero (cross-ROI mean r= 0.024) (**Supplementary Figure 4B**). As with the TTV-corrected analysis, and representative example (split #34) is plotted to (**Supplementary Figure 3D)** illustrate the lack of congruence (r=0.071). Furthermore, there were no regions in humans and mice that showed similar or opposite directionality in effects (**Supplementary Figure 5**).

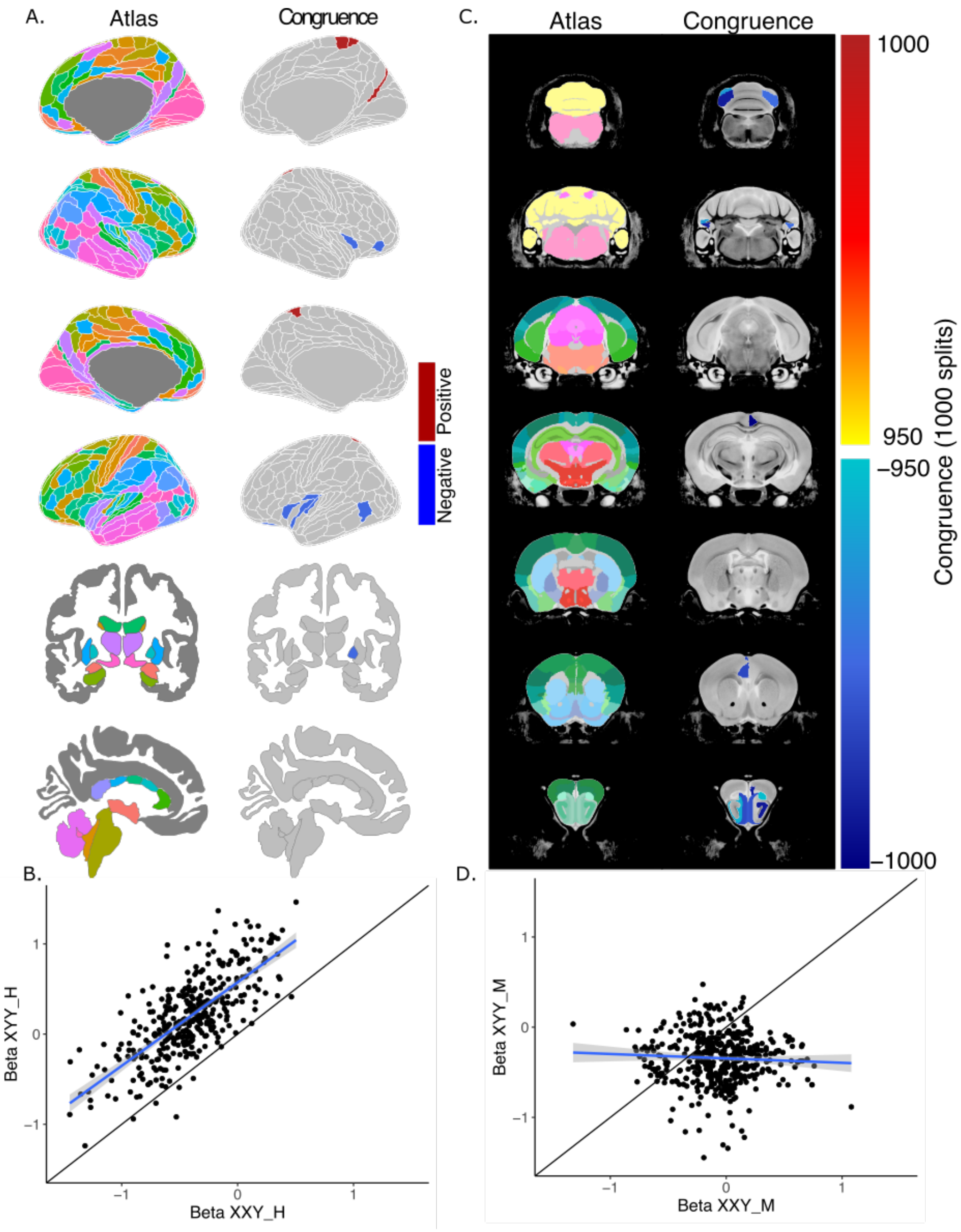


***Supplementary Figure 3. Spatial convergence between added X- or Y- chromosomes on the human and mouse brain without total tissue volume (TTV) correction***. ***A***. *Human brain regions whose volume was either increased (red) or decreased (blue) in both aneuploidy groups.* ***B***. *Correlation between beta values for the XXY_H and XYY_H group across all brain regions.* ***C***. *Mouse brain regions whose volume was consistently increased (red colours) or decreased (blue colours) in 95% (950/1000 splits) of the XY_M control splits in both aneuploidy groups based on bootstrap analysis.* ***D***. *Correlation between beta values for a representative split (34) for XXY_M and XYY_M groups across all brain regions. All plots use the beta values for the effect of aneuploidy with total brain volume correction.*

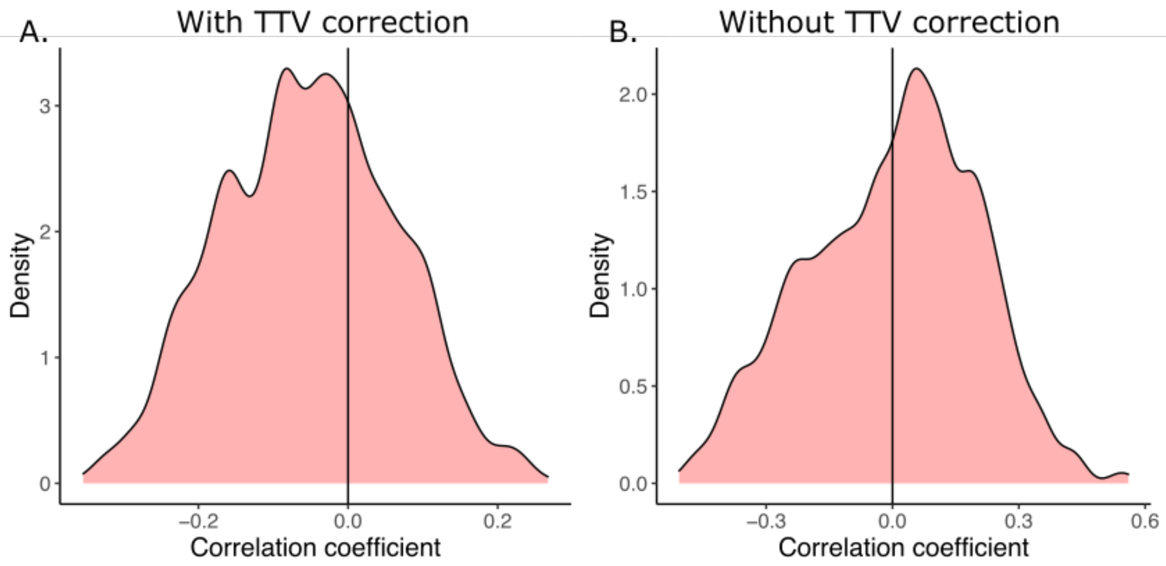


***Supplementary Figure 4. Bootstrap analysis shows low spatial correlation for effects of an added X- or Y-chromosome in the mouse brain.*** *The coefficient for the correlation between cross-ROI standardized effect size (β) for XXY_M (relative to XY_M sample A) and XYY_M (relative to XY_M sample B) both with TTV correction (**A**) and without (**B**).*

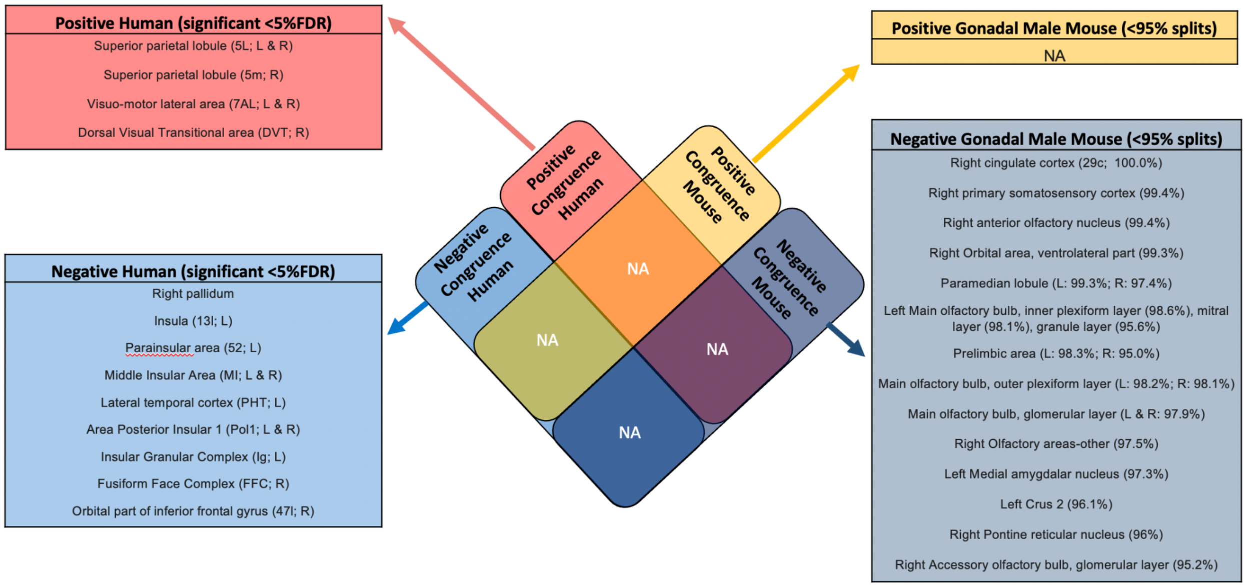


***Supplementary Figure 5. Convergently increased or decreased regions in humans and mice without total tissue volume (TTV) correction.*** *Venn diagram of overlapping ROIs highlighting the regions congruently (for both XXY and XYY) increased or decreased in humans and mice. Brain regions that were statistically significantly impacted by both XXY and XYY in humans alone (<5%FDR) are listed in the red (increased volume) and light blue (decreased volume) boxes on the left. Brain regions that showed significantly convergent impacts of XXY and XYY (based on bootstrap resampling) in mice alone are listed in the yellow (increased volume) and dark blue (decreased volume) boxes. There were no regions showing convergence (increased or decreased), or opposite (increased in one species and decreased in another, or vice versa) highlighted by the "NA" in the intersecting cells.*

#### 1.3 Human-mouse comparison

As explained in **2.4**, the standardized effect sizes for adding an X- or Y- chromosome were compared in the regions identified as human-mouse homologs without correction for TTV. There was low similarity between the effects on the human and mouse brain based on low correlation of standardized effect sizes across regions (added X: r=-0.057; added Y:r=-0.195; **Supplementary Figure 6**). For both humans and mice, an added X-chromosome decreased volume of the right amygdala, bilateral anterior cingulate area, bilateral cerebellum, left nucleus accumbens, right piriform cortex, bilateral primary motor, somatosensory, and retrosplenial cortices, left thalamus, and bilateral ventral orbital area. The left posterior parietal association area, and bilateral primary visual areas were both increased in volume. An added Y-chromosome decreased volume of the left angular insula cortex, bilateral amygdala, bilateral cerebellar cortex, left globus pallidus, left hippocampus, left nucleus accumbens, bilateral piriform cortex, and left temporal association areas, and increased volume in the right hippocampus, right perirhinal cortex, left posterior parietal association area, left primary somatosensory cortex, and left thalamus (**Supplementary Figure 6 & Supplementary Table 1**).

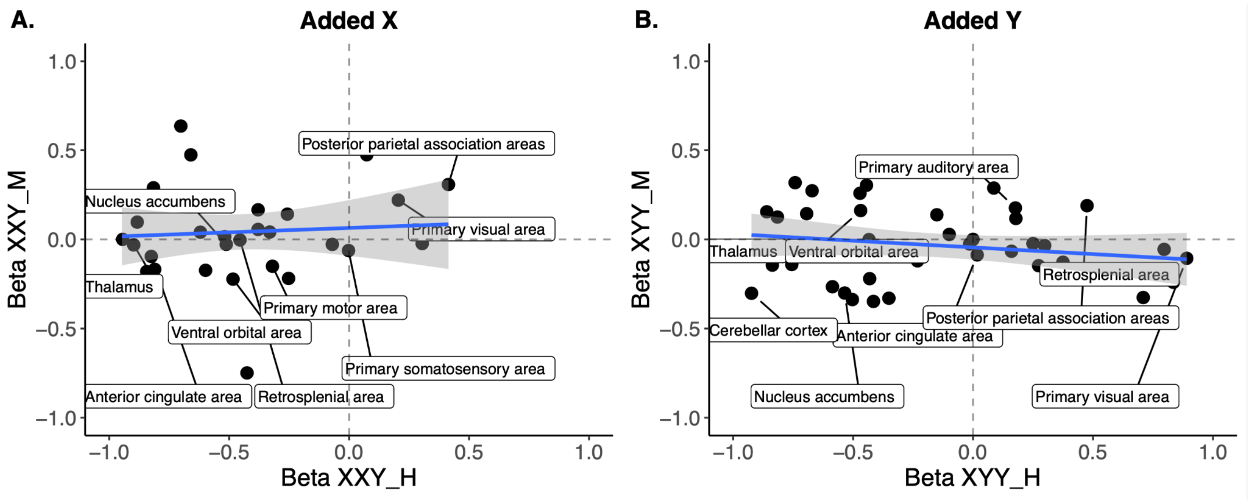


***Supplementary Figure 6. Correlation of effects of added X- or Y- chromosomes in human and mouse homologous brain regions when total tissue volume (TTV) is not controlled for.*** ***A.*** *Standardized effect size correlation for the effect of added X-chromosome (cross-ROI r=-0.06) or* ***B****. added Y-chromosome (cross-ROI r=-0.19). Points are labeled only if they have the same directionality of standardized effect size in humans and mice (i.e., both positive or both negative). For simplicity, labels are provided only for left hemisphere regions given that effects were similar across hemispheres.*

***Supplementary Table 1. Mapping of homologous human-mouse brain regions (standardized effect sizes with TTV correction).*** *Blue shade highlights cells with negative standardized effect sizes, while peach highlights positive ones.*

| | | | | **Beta-value added X** | | | **Beta-value added (Y)** | |
|---|---|---|---|---|---|---|---|---|
| **Label** | **Glasser/Freesurfer* names** | **Mouse atlas** | **Hemisphere** | **Human (XXY_H)** | **Mouse (XXY_M)** | | **Human (XYY_H)** | **Mouse (XYY_M)** |
| **Agranular insula** | AVI, AAIC, MI | Agranular insular area | L | -1.412 | 0.216 | | -0.516 | -0.141 |
| | | | R | -1.503 | 0.299 | | -0.542 | 0.125 |
| **Amygdala** | Amygdala* | Cortical subplate | L | -1.023 | 0.171 | | -0.168 | -0.437 |
| | | | R | -1.147 | -0.017 | | -0.396 | -0.038 |
| **Anterior cingulate area** | A24pr, a24, p24pr, p24, 24dd, 24dv, p32pr, d32, a32pr, p32, s32 | Anterior cingulate area | L | -0.825 | -0.096 | | 0.302 | -0.087 |
| | | | R | -0.810 | -0.168 | | 0.459 | -0.067 |
| **Caudoputamen** | Caudate*, Putamen* | Caudo-putamen | L | -1.185 | 0.241 | | -0.514 | 0.144 |
| | | | R | -1.369 | 0.375 | | -0.504 | 0.273 |
| **Cerebellar cortex** | Cerebellar cortex* | Cerebellar cortex | L | -1.353 | -0.064 | | -0.666 | -0.275 |
| | | | R | -1.444 | -0.195 | | -0.307 | -0.311 |
| **Globus pallidus** | Globus Pallidus* | Pallidum | L | -1.288 | 0.000 | | -0.640 | -0.144 |
| | | | R | -1.270 | 0.116 | | -0.735 | 0.155 |
| **Hippocampus** | Hippocampus* | Hippocampal region | L | -0.883 | 0.113 | | -0.252 | -0.007 |
| | | | R | -0.790 | 0.247 | | 0.052 | 0.132 |
| **Nucleus accumbens** | Nucleus accumbens* | Striatum ventral region | L | -0.512 | -0.054 | | -0.376 | -0.012 |
| | | | R | -0.519 | 0.043 | | -0.037 | 0.023 |
| **Perirhinal area** | PeEc, TF, PHA2, PHA3 | Perirhinal area | L | -0.701 | 0.635 | | 0.449 | -0.024 |
| | | | R | -0.660 | 0.474 | | 0.270 | 0.288 |
| **Piriform cortex** | Pir | Piriform cortex | L | -0.947 | 0.183 | | -0.247 | -0.348 |
| | | | R | -0.844 | -0.180 | | -0.461 | -0.266 |
| **Posterior parietal association areas** | 5m, 5mv, 5L | Posterior parietal association areas | L | 0.414 | 0.366 | | 0.565 | 0.368 |
| | | | R | 0.304 | -0.004 | | 0.863 | -0.031 |
| **Primary auditory area** | A1 | Primary auditory area | L | -0.257 | 0.141 | | 0.342 | 0.176 |
| | | | R | -0.331 | 0.041 | | -0.187 | 0.377 |
| **Primary motor area** | 4 | Primary motor area | L | -0.320 | -0.148 | | 0.115 | -0.030 |
| | | | R | -0.598 | -0.203 | | 0.491 | -0.064 |
| **Primary somatosensory area** | 1, 2, 3a, 3b | Primary somatosensory area | L | -0.002 | -0.063 | | 0.292 | 0.117 |
| | | | R | -0.070 | -0.028 | | 0.897 | -0.056 |
| **Primary visual area** | V1 | Primary visual area | L | 0.205 | 0.220 | | 1.046 | -0.106 |
| | | | R | 0.073 | 0.474 | | 0.995 | -0.241 |
| **Retrosplenial area** | RSC | Retrosplenial area | L | -0.455 | -0.003 | | 0.503 | -0.148 |
| | | | R | -0.252 | -0.219 | | 0.880 | -0.326 |

| | | | | | | | | |
|---|---|---|---|---|---|---|---|---|
| **Subiculum** | PreS | Subiculum | L | -0.379 | 0.171 | | 0.480 | -0.034 |
| | | | R | -0.379 | 0.281 | | -0.218 | 0.270 |
| **Temporal association areas** | FFC, PIT, TE1a, TE1p, TE2a, TF, STV, STSvp, STSva | Temporal association areas | L | -1.086 | 0.246 | | -0.163 | -0.220 |
| | | | R | -0.815 | 0.289 | | -0.177 | 0.304 |
| **Thalamus** | Thalamus* | Thalamus | L | -0.898 | -0.031 | | -0.275 | 0.162 |
| | | | R | -1.045 | 0.023 | | -0.244 | 0.259 |
| **Ventral orbital area** | 10r, 10v | Ventral orbital area | L | -0.484 | -0.223 | | 0.114 | 0.028 |
| | | | R | -0.426 | -0.749 | | 0.581 | -0.416 |
| **Brain stem (midline)** | Brainstem* | Midbrain, Hindbrain | M | -0.620 | 0.040 | | -0.585 | 0.318 |
| **Total tissue volume (midline)** | BrainSegVolNotVent* | | | -0.944 | 0.000 | | 0.423 | 0.000 |
